# Early disruption of entorhinal dopamine in a knock-in model of Alzheimer’s disease

**DOI:** 10.1101/2024.10.10.617678

**Authors:** Tatsuki Nakagawa, Jiayun L. Xie, Marjan Savadkohighodjanaki, Yutian J. Zhang, Heechul Jun, Kai Cao, Ayana Ichii, Jason Y. Lee, Shogo Soma, Yasmeen K. Medhat, Takaomi C. Saido, Kei M Igarashi

**Affiliations:** Department of Anatomy and Neurobiology, School of Medicine, University of California, Irvine; Department of Biomedical Engineering, Samueli School of Engineering, University of California, Irvine; Center for Neural Circuit Mapping, School of Medicine, University of California, Irvine; Center for the Neurobiology of Learning and Memory, University of California, Irvine; Institute for Memory Impairments and Neurological Disorders, University of California, Irvine; Lab for Proteolytic Neuroscience, RIKEN Center for Brain Science, Japan

## Abstract

The entorhinal cortex (EC) is a critical brain area for memory formation, while also the region exhibiting the earliest histological and functional alterations in Alzheimer’s disease (AD). The EC thus has been long hypothesized as one of the originating brain areas of AD pathophysiology, although circuit mechanisms causing its selective vulnerability remain poorly understood. We found that dopamine neurons projecting their axons to the lateral EC (LEC), critical for memory formation in healthy brains, become dysfunctional and cause memory impairments in early AD brains. In amyloid precursor protein knock-in mice with associative memory impairment, LEC dopamine activity and associative memory encoding of LEC layer 2/3 neurons were disrupted in parallel from the early pathological stage. Optogenetic reactivation of LEC dopamine fibers, as well as L-DOPA treatment, rescued associative learning behavior. These results suggest that dysfunction of LEC-projecting dopamine neurons underlies memory impairment in AD from early stages, pointing to a need for clinical investigation of LEC dopamine in AD patients.

## Introduction

The memory circuit consisting of the entorhinal cortex and hippocampus is critically involved in memory formation and retrieval, and damage to this circuit will lead to the impairment of memory formation^1,2^. The entorhinal cortex has been hypothesized as one of the most plausible originating brain regions of AD pathophysiology^3–5^. Past histological studies from post-mortem AD brains have established that the entorhinal cortex has atrophies earlier than the hippocampus, with entorhinal cortex layer 2 neurons being the most vulnerable from the early disease stage^3,6,7^. A subsequent functional magnetic resonance imaging (fMRI) study from early-stage AD patients showed that, among the entorhinal-hippocampal circuit, the lateral entorhinal cortex (LEC) suffers the earliest loss of overall activity, earlier than the medial entorhinal cortex (MEC)^8^. However, it remains unclear why pathophysiological abnormalities of AD appear earliest in the LEC, and what type of brain function is lost from the LEC in AD. In healthy brains, the LEC receives sensory information from olfactory and somatosensory areas^9–11^, exhibiting firing to surrounding objects and items that have olfactory, somatosensory, and visual features^12-^_15_. Using an olfactory item-outcome associative memory task, we recently identified in healthy mice that LEC layer 2a (LEC_L2a_) fan cells represent novel odor cues associated with reward outcome, and this representation is controlled by dopamine inputs to the LEC from the ventral tegmental area (VTA) and substantia nigra pars compacta (SNc)^16^. Because dopamine neurons are known to be intrinsically vulnerable to aging due to their high metabolic stress and elevated mitochondrial oxidant^17–19^, we asked if LEC-projecting dopamine neurons underlies memory deficit symptoms in AD.

## Results

### Impaired associative memory formation in APP-KI mice

To investigate the LEC in AD brains, we used amyloid precursor protein knock-in (APP-KI) mice^20,21^. The APP-KI mouse recapitulates the gradual process of AD pathological progression more accurately than aggressive transgenic AD mouse models. In these mice, age-dependent accumulation of Aβ starts widely in cortical regions, including the LEC, at 2 months of age (mo) (**Fig. 1a and Extended Data** Fig. 1). We tested mice’s memory formation using an LEC-dependent olfactory cue-outcome associative memory task^16^ (**Fig. 1b**). In this task, mice use pre-learned knowledge to rapidly form associative memories between novel odor cues and their outcomes. Mice were initially trained with pre-learning sessions, where they learned to lick after Odor-A for sucrose water reward, and to withhold licking after Odor-B to avoid quinine water punishment. After mice learned to discriminate at >80% correct performance, daily associative learning sessions were tested, where Odors-A, -B, and a pair of novel odors (Odor-1 → sucrose, Odor-2 → quinine) were randomly presented. Associative learning was tested daily in individual mice using novel rewarded odors (Odors-C, -E, -G, …; collectively termed Odor-1) and novel punished odors (Odors-D, -F, -H, …; termed Odor- 2). This associative learning was previously shown to be dependent on the activity of LEC layer 2a neurons as well as the dopamine inputs to the LEC^16^.

**Figure 1.**
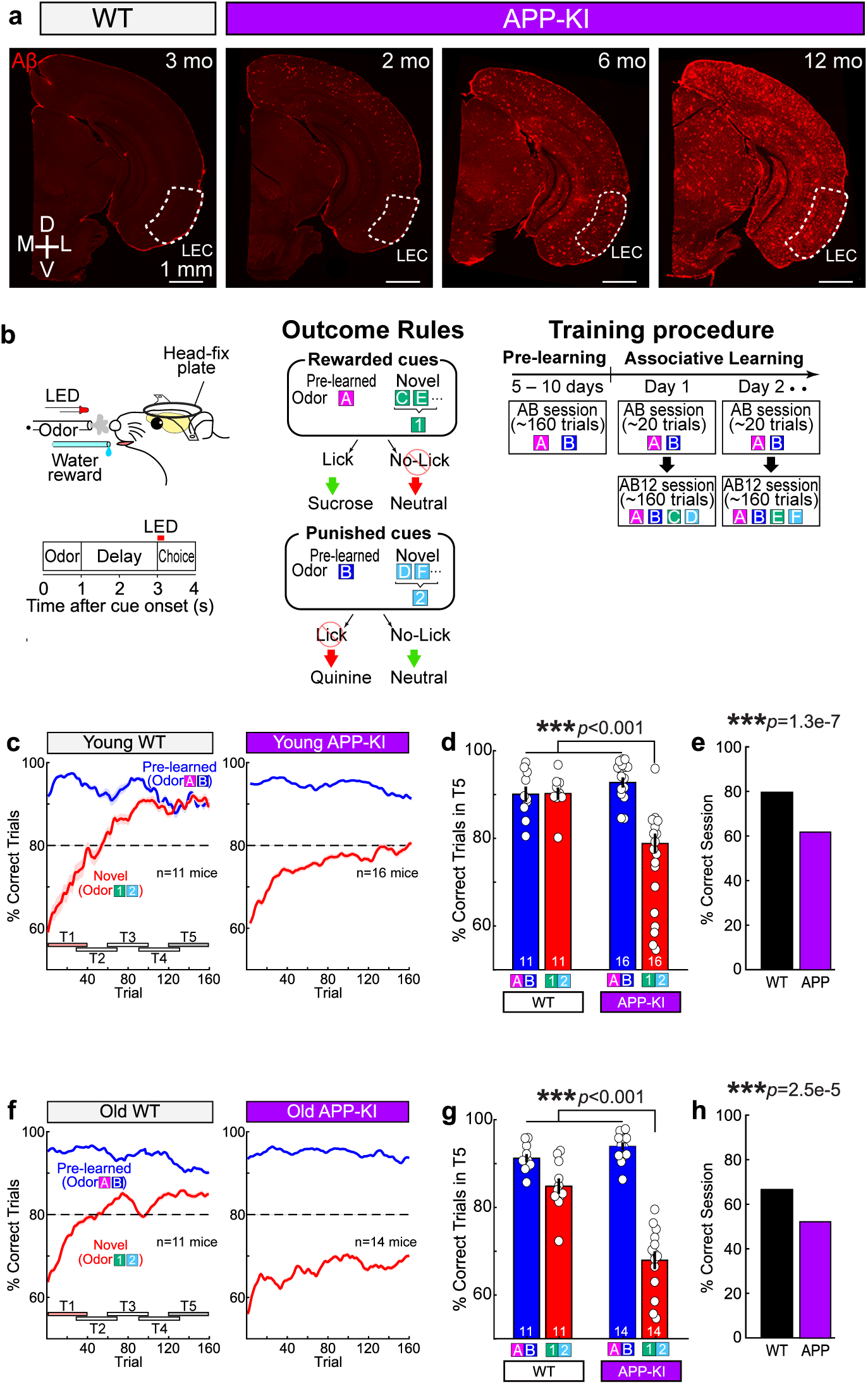
Early impairment of associative memory formation in APP-KI mice **(a)** Coronal section with anti-Aβ immunostaining for 3-month-old WT and 2, 6, 12- month-old APP-KI mice. D, dorsal, V, ventral, M, medial, L, lateral. **(b)** Head-fixed mice learned associations between odor cues and licking for sucrose water reward. During associative learning sessions, animals were tested with AB-only and AB12 sessions with novel odors (C/D, E/F, or G/H …). Novel odors are collectively referred to as Odor-1 and Odor-2 in subsequent figures. **(c)** Correct trial rate for pre-learned odors (A/B, blue) and novel odors (1/2, red) in young WT mice (3-6 mo, left) and young APP-KI mice (3-6 mo, right). Timepoints T1-T5 (rectangles) are shown. **(d)** Percentage of correct trials in T5 in young WT (n = 11) vs young APP-KI (n = 16) mice (p=0.051, ANOVA; p<0.001, post-hoc Tukey test). **(e)** Percentage of sessions where mice correctly learned new association (p=1.3e- 7, binomial test). **(f)** Same as Fig. 1c, but for old WT (7-11 mo, left) and old APP-KI mice (7-11 mo, right). **(g)** Same as Fig. 1d, but for old WT (n = 11) and old APP-KI (n = 14) mice (p= 2.4e- 5, ANOVA; p<0.001, post-hoc Tukey test). **(h)** Same as Fig. 1e, but for old WT and old APP-KI mice (p=2.5e-5, binomial test).

Because the APP-KI mouse is on the C57BL/6 background, we used C57BL/6 mice as a control (referred to as WT mice). Consistent with our previous result, control 4-mo WT mice (defined as young WT mice) rapidly learned new associations by trial and error within a 160-trial session (red trace, **Fig. 1c**, Left), while continuously retrieving pre-learned Odors-A/B associations with >80% correct rate throughout a session (blue trace, **Fig. 1c**, Left, **and 1d**). By contrast, 3-6 mo APP-KI mice (young APP-KI mice) showed decreased and slower learning performance for new associations, barely reaching the 80% criteria (**Fig. 1c**, Right**, and 1d; Extended Data** Fig. 2). We divided the session into five timepoints^16^ (T1-T5, **Fig. 1c**) and compared the new associations performance between WT and APP-KI mice in T5 (**Fig. 1d**). Young WT mice performed at 90.6 ± 0.9% in T5, whereas young APP-KI performed at 78.6 ± 2.1% (p=4.1e-5 ANOVA; p=3.1e-8, post-hoc Tukey test; n = 11 young WT and n = 16 young APP-KI mice). We defined sessions with >=80% performance for new associations as correct sessions, and sessions with <80% performance as error sessions. Young WT mice had correct sessions at 79.6% among all tested sessions, whereas young APP-KI had correct sessions at 61.4% (p=1.3e-7, binomial test; **Fig. 1e**). At older age point of 7-11 mo, APP-KI mice (defined as old APP-KI mice) showed further impairment in associative memory formation compared to age-matched old WT mice (84.9 ± 1.8% for n = 11 old WT mice vs. 67.9 ± 2.0% for n = 14 old APP-KI mice; p=5.4e-14 ANOVA; p=2.7e-9, post-hoc Tukey test; **Fig. 1f and 1g**). Notably, old WT mice showed a slight decline of associative memory compared to young WT mice (**Extended Data** Fig. 2c, p=0.031 ANOVA; p=0.038, post-hoc Tukey test; n = 11 young WT and n = 11 old WT mice). Old WT mice correctly learned 67.0% of all sessions, whereas APP-KI correctly learned 52.3% (p=2.5e-5, binomial test; **Fig. 1h**). By contrast, both young and old APP-KI mice showed intact performance for retrieving pre-learned associative memories (Odors-A/B, blue traces in **Fig. 1c and 1f**; p>0.05, **Fig. 1d and 1g**), suggesting that olfactory perception and memory retrieval are both spared in APP-KI mice. APP-KI and WT mice showed comparable duration for the training with pre-learning sessions using Odor-A and Odor-B (p=0.13 ANOVA; post-hoc Tukey test p=0.99 between young APP-KI vs young WT mice, p=0.91 between old APP-KI vs old WT mice**; Extended Data** Fig. 2f). Together, these results demonstrate that APP-KI mice show progressive impairment in the formation, but not the retrieval, of associative memory starting at 4 mo.

### Disrupted associative memory encoding in LEC_L2/3_ neurons

We next investigated memory-encoding spike activity of LEC neurons in AD brains. We implanted recording drives with 64 electrodes targeting LEC layers 2/3 (LEC_L2/3_) and recorded 479 and 846 neurons from n=5 young WT and n=5 young APP-KI mice, respectively (**Fig. 2a and Extended Data** Fig. 3). LEC_L2/3_ neurons in APP-KI mice showed decreasing spike widths compared to those of WT mice, suggesting alterations in spike generation mechanisms (**Extended Data** Fig. 4a-b). Consistent with Aβ-induced neuronal hyperactivity observed previously in the hippocampus^22,23^, the mean firing rates of putative LEC_L2/3_ principal neurons and interneurons were both higher in APP-KI mice than those of WT mice (**Extended Data** Fig. 4c). An example WT neuron from a correct session (**Fig. 2b**) responded to pre-learned rewarded Odor-A from the initial 10 trials (T1, black asterisks in **Fig. 2b**; p<0.01, rank sum test compared to pre-cue period). This cell quickly acquired a response also to novel rewarded Odor-1, and these responses continued through the last 10 trials (T5, p<0.01, red asterisks in **Fig. 2b**). Thus, this neuron underwent a plastic change from an “Odor-A cell” at T1 to an “Odor-A/1 cell” at T5 during learning. By contrast, an example APP-KI neuron from an error session (**Fig. 2c**) showed spike firing only to Odor-A through T1 to T5. We analyzed response properties of the LEC_L2/3_ population. While 16% of LEC_L2/3_ neurons fired to at least one odor cue at T5 in WT mice, an excessive 34% of neurons exhibited cue responses in APP-KI mice (**Fig. 2d**. p=3.6e-9, chi square test; p<0.01 for 1-odor and 2-odor responsive cells, Benjamini-Hochberg post-hoc test). Among these cue-responsive neurons in APP-KI mice, Odor-A cells and Odor-A/1 cells were constantly greater through T1-T5 compared to those in WT mice (**Extended Data** Fig. 5, p<0.001, chi square test; p<0.05, Benjamini-Hochberg post-hoc test; see **Extended Data** Fig. 6 for cell responses). Population representations were then assessed using principal component analysis^16^ (PCA, **Fig. 2e-f, Extended Data** Fig. 7). For this analysis, we used all sessions including both correct and error sessions to assess overall alteration (see **Extended Data** Fig. 8 for separate PCA for correct and error sessions). In WT mice, trajectories of Odors-A and -1 stayed close in principal component (PC) space (**Fig. 2e**). These similar representations between Odors-A and -1 are the characteristic property for generalizing rewarded cues found previously in LEC layer 2a fan cells^16^. By contrast, in APP-KI mice, trajectories for the individual four cues remained far apart throughout the session, separating representations of Odor-A from Odor-1 (**Fig. 2f**). When sessions were separated between correct and error sessions, the PC trajectories for Odors-A and -1 were insufficiently close in correct sessions, while they were completely separated in error sessions (**Extended Data** Fig. 8). We measured Euclidian distances between trajectories to assess similarity of spike representations between cue types (**Fig. 2g**). The ninety-fifth percentile distance obtained from shuffled data was used as a threshold for statistical significance (red line, **Fig. 2g**). In WT mice, A-1 distance became smaller than the shuffle through T2 to T5, indicating generalized representations of rewarded cues during learning. However, A-1 distance in APP-KI mice remained above shuffle, indicating that Odors-A and -1 representations were not generalized together in APP-KI mice. This dissimilarity is apparent using a Similarity Index (SI)^16^ (see Method, **Fig. 2h**). A bootstrapping analysis confirmed the decreased A-1 similarity in APP-KI mice compared to WT mice throughout the session (p<0.001 in T1 – T5, **Fig. 2i**). In old age, generalization of Odors-A and -1 was observed through T2-T5 in WT mice, whereas representations for these two cues were kept separated throughout the session again in APP-KI mice (**Extended Data** Fig. 9-10). Together, these data demonstrate that the spike representation for generalizing Odor-A and Odor-1, observed in healthy WT mice, was disrupted in APP-KI mice from a young age.

**Figure 2.**
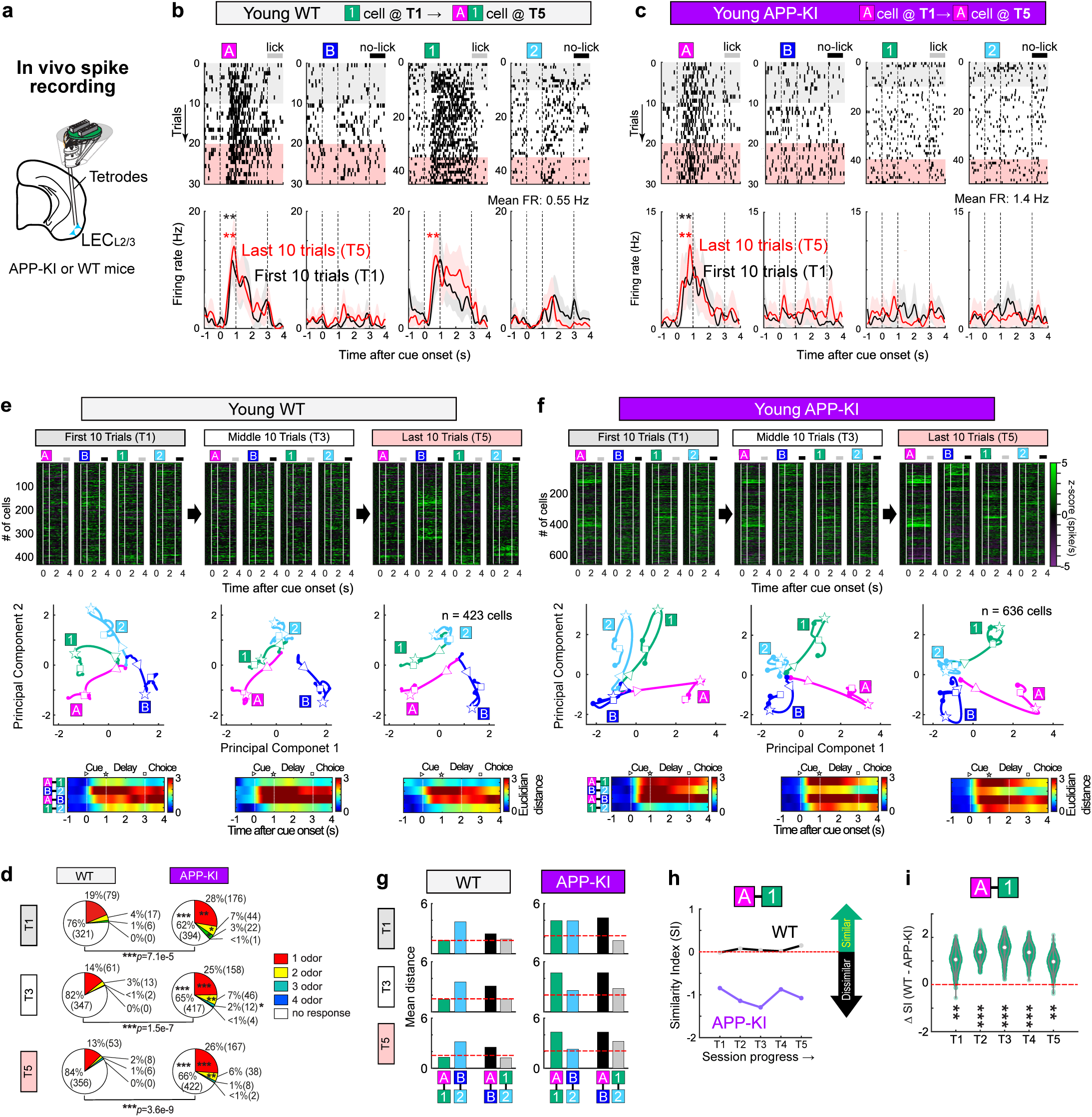
Disrupted cue-reward association of LEC_L2/3_ neurons in young APP-KI mice (a) A schematic diagram of the in vivo spike recording experiment. (b) An example LEC_L2/3_ cell in a young WT mouse from a correct session, showing firing for Odor-A and developing firing for Odor-1. (**p<0.01 during cue-delay period compared to pre-cue period, Wilcoxon signed-rank test). Black trace, first 10 trials. Red trace, last 10 trials. (c) An example LEC_L2/3_ cell in a young APP-KI mouse from an error session exhibited constant firing for Odor-A with no change in its firing pattern. (d) Percentage of LEC_L2/3_ cells classified by odor cue response in the first 10 trials (T1, top), middle 10 trials (T3, middle), and last 10 trials (T5, bottom) (***p<0.001, Chi-squared test for distributions between WT and APP-KI mice; *p<0.05, **p<0.01, ***p<0.001, Benjamini-Hochberg post-hoc test for each responsive class between WT and APP-KI mice; n = 423 cells from young WT and n= 636 cells from young APP-KI mice). Percentages of neurons in each responsive class is shown, and numbers in parentheses denote number of neurons in each responsive class. (e) (Top) Spike firing rate of n = 423 LEC_L2/3_ cells from young WT mice, shown in z- score during first 10 (T1), middle 10 (T3) and last 10 (T5) trials the session. (Middle) Time-resolved principal component analysis (PCA) trajectory of LEC_L2/3_ cell activity for each odor type (▷ cue-onset; ⋆ cue-offset/delay-onset; □ delay- offset). (Bottom) Euclidian distance between odor types. (f) Same as Fig. 3e, but for young APP-KI mice (n = 636 cells). Trajectories for Odor- A and Odor-1 kept far apart through T1 to T5. (g) Mean Euclidian distance between PCA trajectories during 0.5 – 1.5 s after cue onset in young WT (left) and young APP-KI mice (right). Ninety-fifth percentile distance obtained from shuffled data denotes significant distance (red line). (h) Similarity index (SI) between Odor-A and -1 in young WT and young APP-KI mice across learning. Positive SI denotes similar representations between Odor-A and Odor-1, whereas negative SI denotes dissimilar representations. (i) Similarity Index (SI) of young WT and young APP-KI mice was compared using bootstrapping method (see Method). SI was calculated for 1000 bootstraps, then SIs for WT were subtracted by SIs for APP-KI. The subtraction confirms higher similarity in the representations between Odor-A and Odor-1 in WT mice compared to that in APP-KI mice (**p<0.01, ***p<0.001; n=1000 bootstrapping test).

### Dysfunction of LEC dopamine activity

In healthy brains, the spike encoding of generalizing rewarded cues in LEC neurons is controlled by dopamine inputs from the ventral tegmental areas (VTA) and substantia nigra pars compacta (SNc) to the LEC^16^. Our previous optogenetic inhibition of LEC dopamine in healthy mice led to impaired associative memory encoding of LEC L2a neurons as well as animals’ associative learning^16^, similar to the results obtained above in APP-KI mice. Thus, we next investigated LEC dopamine activity in APP-KI mice. Because APP-KI mice already exhibit disruption of LEC spike activity from young age of 3-6 mo, we focused our analyses on this early stage. Aβ accumulation emerged from 4 mo in midbrain regions including the VTA and SNc (**Fig. 1**). To assess whether dopamine axons degenerate in the LEC, we performed immunohistochemistry for tyrosine hydroxylase (TH) and measured the density of TH^+^ axons in the LEC (**Fig. 3a-c**). Although there was a trend of decrease in the density of TH^+^ axons of APP-KI mice, the effect was not significant across genotype nor age (**Fig. 3c**, p=0.38 between APP-KI vs WT, p=0.46 between old vs young; two-way ANOVA). We also measured a density of dopaminergic neurons in the VTA and SNc, finding no significant alteration between APP-KI and WT mice (**Extended Data** Fig. 11).

**Figure 3.**
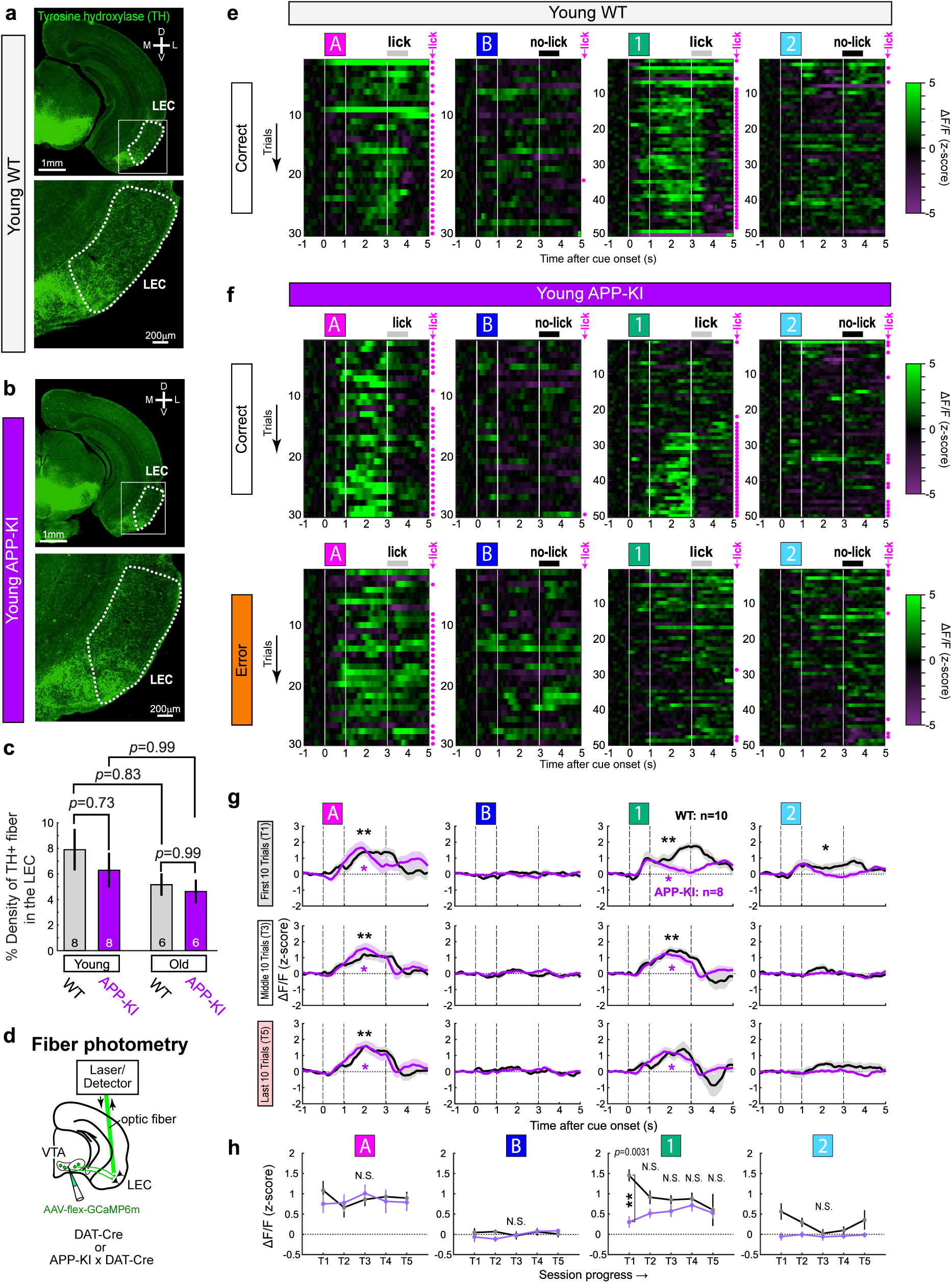
Dysfunction of LEC dopamine in young APP-KI mice **(a)** Coronal section with anti-tyrosine hydroxylase (TH) immunostaining for young WT mice. D, dorsal, V, ventral, M, medial, L, lateral. **(b)** Same as Fig. 3a, but for young APP-KI mice. **(c)** Density of TH-positive fibers in the LEC of WT and APP-KI mice (p= 0.38, ANOVA). P-values for post-hoc Tukey test are shown in the figure (n = 8 young WT mice, n = 8 young APP-KI mice, n = 6 old WT mice, n = 6 old APP-KI mice). All post-hoc p-values, including those not shown in the figure, were above 0.05. **(d)** Fiber photometry recording of calcium signals from dopamine fibers in the LEC. **(e)** Example recording of GCaMP signals from LEC dopamine fibers shown in z- scored ΔF/F, from a young WT mouse during a correct session. **(f)** Same as Fig. 3e, but for young APP-KI mice during correct (Top) and error (Bottom) sessions. **(g)** GCaMP signals in the first 10 trials (T1, top), middle 10 trials (T3, middle), and last 10 trials (T5, bottom). Photometry signals from both correct and error sessions were averaged for WT and APP-KI mice. *p<0.05, **p<0.01, ***p<0.001, signals during 1-4 s after cue onset compared with those during 1 s pre-cue period, Wilcoxon signed-rank test; n = 10 and n = 8 hemispheres from WT and APP-KI mice, respectively. **(h)** Mean GCaMP signals in APP-KI mice showed decreasing signals on Odor-1 trials compared to WT mice (p= 0.0013, ANOVA; p= 0.0031 in T1, post-hoc Tukey test; n = 10 and n = 8 hemispheres from WT and APP-KI mice, respectively).

Because LEC dopamine fibers did not show significant morphological degeneration in APP-KI mice, we next examined functional activity of LEC dopamine axons using fiber photometry^16^. Adeno-associated virus (AAV) expressing calcium activity indicator GCaMP6m in a Cre-dependent manner (AAV-flex-GCaMP6m) was injected in the VTA/SNc of young APP-KI mice crossed with DAT-Cre (APP-KI x DAT-Cre), and photometry fibers were implanted in the LEC (**Fig. 3d**). In control DAT-Cre mice, LEC dopamine supplied reward expectation signals during rewarded trials as observed previously^16^ (**Fig. 3e** and **g**; Extended Data Fig. 12a and 12c. At T1, dopamine was released primarily during delay periods of pre-learned rewarded Odor-A (mean z-scored GCaMP signal, 1.1 ± 0.24; p=1.8e-4, rank sum test compared to pre-cue period). The signal also emerged for novel Odor-1 (1.8 ± 0.2; p=1.8e-4) and Odor-2 cues (0.76 ± 0.24; p=0.025). Subsequently, dopamine signals disappeared for Odor-2, while remaining present for rewarded Odor-A and Odor-1 through T5. By contrast, when APP-KI mice managed to slowly learn the association (correct session, **Fig. 3f**, top**; Extended Data** Fig. 12b-c), dopamine signals for Odor-A remained intact (p=0.66, ANOVA, **Fig. 3h**), while signals for Odor-1 emerged in a delayed manner from T3 (p=0.0013, ANOVA; p=0.0031 in T1, post-hoc Tukey test, **Fig. 3h**). The delayed emergence of Odor-1 dopamine signals correlated well with slow learning of licking to Odor-1 in APP-KI mice (*r*(19) = 0.78, p=2.5e-05, Pearson correlation, **Extended Data** Fig. 12d), implying that slow emergence of LEC dopamine signals causes delayed learning in APP-KI mice. When animals did not learn associations (error sessions), dopamine signals for Odor-1 were completely absent, while the signals for Odor-A again remained present (**Fig. 3f**, bottom and **Extended Data** Fig. 12b-c). Together, these data demonstrate that, while the LEC dopamine fiber structure was relatively intact, LEC dopamine activity for novel rewarded cues became dysfunctional from a young age in APP-KI mice.

### Rescued memory by dopamine reactivation

Having established that LEC dopamine is diminished in young APP-KI mice, we finally asked whether the reactivation of LEC dopamine could rescue associative memory in APP-KI mice. First, to directly reactivate LEC dopamine, we optogenetically stimulated LEC dopamine fibers while young APP-KI mice performed the associative memory task (**Fig. 4a**). Channelrhodopsin-2 (ChR2) was expressed in dopamine fibers of APP-KI x DAT-Cre mice using AAV-DIO-ChR2 injected in the VTA/SNc, and optic fibers for laser stimulation were bilaterally implanted above the LEC. Because LEC dopamine was diminished largely during novel rewarded Odor-1 trials (**Fig. 3**), we stimulated LEC dopamine fibers only on Odor-1 trials (**Fig. 4a**). When LEC dopamine axons were stimulated, APP-KI mice showed significantly enhanced performance for the acquisition of novel odor associations compared to no-stimulation control sessions (stimulation: 95.2 ± 1.6% vs. control: 77.9 ± 4.9% in T5; p=0.035, ANOVA; p=0.024, post-hoc Tukey test; n = 11 young APP-KI mice; **Fig. 4b-c and Extended Data** Fig. 13a). The percentage of total correct sessions significantly increased with stimulation (stimulation 84.4% vs. control 51.1%; p=7.1e-6, binomial test; **Fig. 4d**).

**Figure 4.**
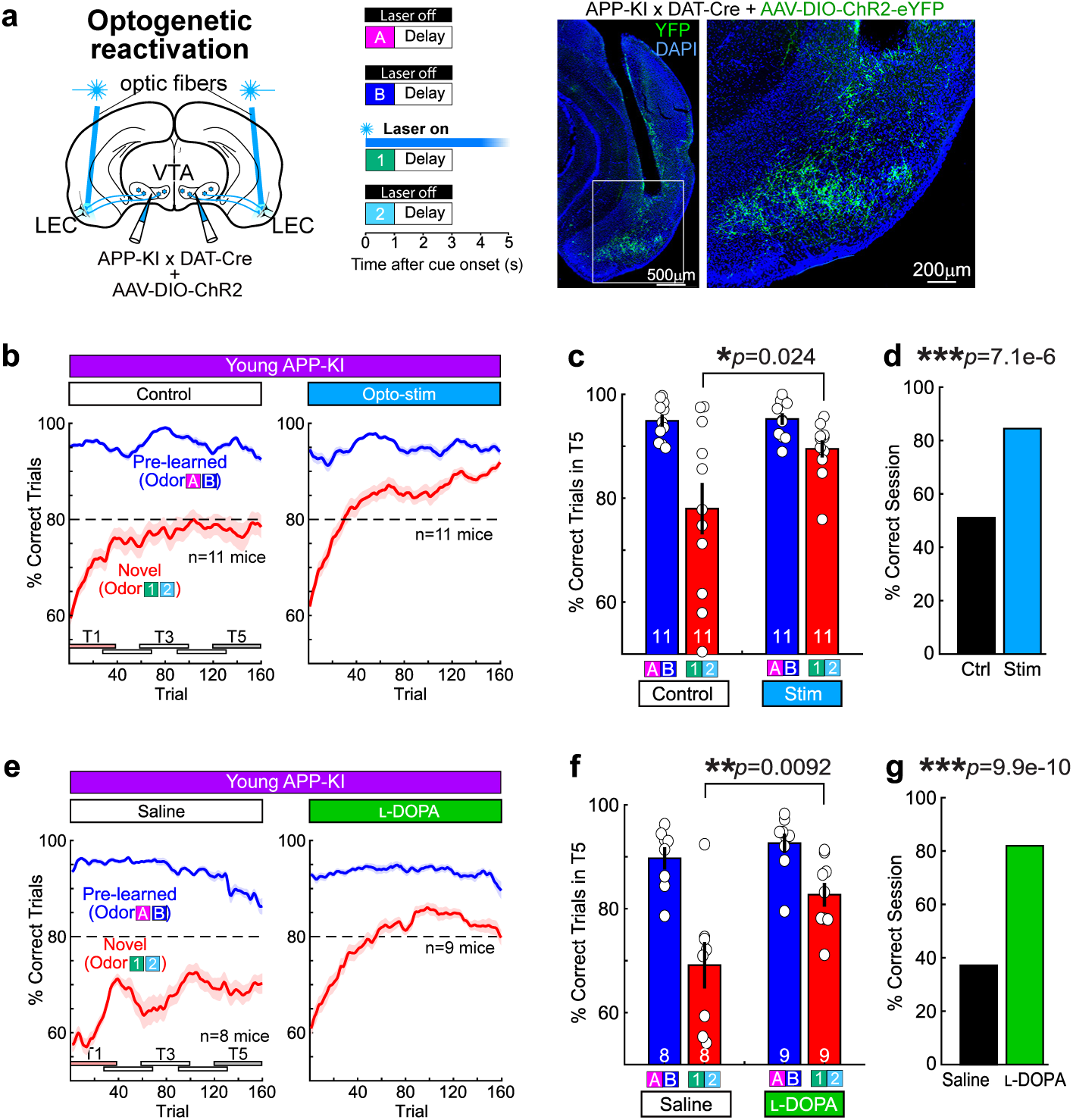
Reactivation of LEC dopamine rescued associative memory of APP-KI mice **(a)** (Left) Optogenetic stimulation of dopamine fibers in the LEC during Odor-1 trials. (Right) A section from stimulated mouse showing an optic fiber track and ChR2- expressing dopamine fibers in the LEC. **(b)** Correct trial rate for pre-learned odors (A/B, blue) and novel odors (1/2, red) in control no-stimulation sessions (left) and optogenetic stimulation sessions (right), obtained from n=11 young APP-KI mice. **(c)** Percentage of correct trials in T5 of control sessions and optogenetic stimulation sessions (p= 0.035, ANOVA; p= 0.024 for Odor-1/2 between control sessions vs. stimulation sessions, post-hoc Tukey test; n = 11 young APP-KI mice). **(d)** Percentage of sessions where mice correctly learned new association (p=7.1e-6, binomial test). **(e)** Correct trial rate for pre-learned odors (A/B, blue) and novel odors (1/2, red) in saline-injected young APP-KI mice (n=8, left) and L-DOPA-injected young APP-KI mice (n=9, right) **(f)** Percentage of correct trials in T5 in saline-injected young APP-KI mice and L-DOPA-injected young APP-KI mice (p= 0.0063, ANOVA; p= 0.0092 for Odor-1/2 in Saline vs. Odor-1/2 in L-DOPA treatment, post-hoc Tukey test; n = 8 saline mice and n=9 L-DOPA mice). Percentage of sessions where mice correctly learned new association (p=9.9e-10, binomial test).

We finally tested L-DOPA treatment, a precursor of dopamine used for the treatment of Parkinson’s disease patients^24^. Young APP-KI mice injected intraperitoneally with L- DOPA recovered their associative memory up to the level of WT mice, with enhanced performance compared to saline injected groups (L-DOPA groups 82.6 ± 2.3% vs. saline group 69.1 ± 4.5% at T5; p=0.0063, ANOVA; p=0.0092, post-hoc Tukey test; n = 9 L- DOPA mice, n = 8 saline mice; **Fig. 4e-f and Extended Data** Fig. 13b). The percentage of total correct sessions significantly increased with L-DOPA treatment (L-DOPA 82.6%, saline 69.1%; p=9.9e-10, binomial test; **Fig. 4g**). Together, these data indicate that LEC dopamine dysfunction underlies the associative memory impairment in young APP-KI mice. Our results further suggest that reactivation of LEC dopamine has the potential to rescue associative memory formation in AD-impacted brains.

## Discussion

Although the role of the entorhinal cortex, especially the LEC, has been long hypothesized in AD pathogenesis, it has remained unclear why this region suffers selective dysfunction. Here we obtained solid circuit-level evidence that dopamine inputs from the VTA/SNc to the LEC become dysfunctional in the early stage of AD brains, causing associative memory impairments. Although the pathogenic role of dopamine neurons in Parkinson’s disease has been well documented, its role in AD has remained poorly understood due to obscurity of dopamine projections to the entorhinal-hippocampal memory circuit. Among this circuit, the hippocampus receives limited VTA/SNc axons, but rather receives axons from the locus coeruleus that co-release dopamine and norepinephrine^25,26^. Our recent identification of LEC dopamine in healthy brains^16^ paves a way to investigate its role in AD pathogenesis. We propose that the selective functional vulnerability of LEC in AD is attributed, at least partially, to the dopamine projection pattern specific to the LEC among the entorhinal-hippocampal circuit. Dopamine fibers have been identified not only in the LEC of rodents^27,28^ but also in the entorhinal cortex of humans^29–33^, while no research to assess entorhinal dopamine functions has been performed in AD patients. Our results highlight the need to investigate entorhinal dopamine functions in human AD brains.

In young APP-KI mice at 4 mo, the density of LEC dopamine fibers was comparable to that of age-matched WT mice, suggesting limited axonal degeneration at the early AD stage (**Fig. 3**). By contrast, dopamine signals for novel rewarded Odor-1 were significantly impaired in APP-KI mice, while signals for familiar rewarded Odor-A were relatively intact. These results suggest that, in young APP-KI mice, dopamine reserves are available at synaptic terminals in the LEC, yet fail to be released for novel rewarded Odor-1 cues. Our result of optogenetically-rescued associative memory also supports spared dopamine reserves in APP-KI mice. Because photometry detects calcium events induced by spiking activity in axon terminals, it is likely that Aβ damages either the spike generation mechanism in the soma of LEC-projecting dopamine neurons located in the VTA/SNc, or axonal spike propagation mechanisms toward the LEC. The decreased dopamine release for novel rewarded Odor-1 impaired memory encoding of LEC neurons, presumably due to the lack of dopamine-dependent neuronal plasticity^34,35^. Future work is required to decipher the cellular mechanism of how Aβ accumulation leads to LEC dopamine dysfunction.

Aβ accumulation may also have impaired intrinsic spike mechanisms in the LEC circuit. The hyperactivity of both LEC_L2/3_ principal neurons and interneurons, as well as the elevated number of neurons firing to odor cues (**Fig. 2**) imply abnormality in the E-I balance. The excessive numbers of cue-responsive neurons presumably decreased the signal-to-noise ratio of LEC_L2/3_ representations, which, together with diminished LEC dopamine, blocked the generalization of Odor-A and Odor-1. Even with such alterations of LEC_L2/3_ neurons, the reactivation of LEC dopamine fibers could improve associative memory formation of APP-KI mice (**Fig. 4**). We presume that stimulating dopamine release during novel reward cues reactivates plasticity of LEC_L2/3_ neurons, rescuing the generalization of reward cues. Although optogenetic stimulation is not extendable to the treatment of AD patients, L-DOPA treatment is widely used to increase global dopamine levels in Parkinson’s disease patients. Our results raise the possibility that L-DOPA could also relieve memory impairments in early-stage AD symptoms. The mechanistic understanding of LEC dopamine dysfunction is expected to open a pathway to a future treatment for ameliorating memory functions in AD patients.

## Acknowledgments

We thank Dr. Masashi Kitazawa and members in the Igarashi lab for providing valuable comments on the work.

## Author contributions

K.M.I., S.S and T.N. conceived the project and designed the experiments. T.N., J.L.X., Y.J.Z., M.S., H.J., J.Y.L., and S.S. performed the experiments.

T.N. performed the analyses. T.C.S generated and provided the APP-KI mice. T.N., J.Y.L., and K.M.I. wrote the paper with input from all authors.

## Data, code and materials availability

Neurophysiological data and analytical codes are available upon request, and will be deposited with a subsequent protocol paper.

Supplementary materials contain Methods, Extended Data Figures 1-13 and Extended Data Tables 1-4.

## Methods

### Subjects

All procedures were conducted in accordance with the guidelines of the National Institutes of Health and approved by the Institutional Animal Care and Use Committee at the University of California, Irvine. Mice were maintained in standard housing conditions on a reversed 12h dark/light cycle with food and water provided *ad libitum.* All experiments were conducted during the dark phase. Mouse lines and their received procedure are summarized in **Extended Data Table 2 and 4**.

All animals had either C57BL/6 background or were backcrossed to C57BL/6 for at least 7 generations. They were housed individually with an exercise wheel following their first procedure. Medications and appropriate treatments were applied across 1-2 weeks for recovery. If animals died before the post-experiment histological analysis, their recording positions could not be validated and therefore were removed from the data analysis. Similar numbers of males and females were used in the experiments (see **Extended Data Table 2**).

### Electrode, Drive and Optic Fiber Preparation

*Tetrode drive.* Custom-built 64-channel drives containing bundled tetrodes were used for recording as previously described (Lee et al., 2021).

*Optic fibers*. For optogenetic inhibition experiments, optic fibers of 400 µm diameter and 10-13mm length (Thorlabs) were used.

### Surgery

All surgical procedures were performed as previously described (Lee et al., 2021). Mice received post-operative injections of flunixin and enrofloxacin daily as needed.

*Injections.* For injections targeting the VTA/SNc, a 1 mm craniotomy was centered at AP 3.15 mm, ML 0.8 mm from bregma. The micropipette was lowered to a depth of 4,000 µm from the brain surface. Viruses were manually injected at approximately 0.1 µL/min using a hydraulic manipulator (MO-10, Narishige). AAVs used in the study are summarized in **Extended Data Table 4**.

*Tetrode drive and optic fiber implantations.* After allowing at least one week recovery from injection surgeries, mice were implanted with tetrode drives or optic fibers as previously described (Lee et al., 2021). Briefly, a custom titanium head-plate was affixed to the skull, then craniotomies were drilled at the LEC coordinates (AP 3.5 mm, ML 3.5 mm from bregma, depth 3300-3500 μm from the brain surface, 10 degrees lateral). For drive implants, reference wire was implanted on the cerebellar surface. Drives or optic fibers were secured to the skull with dental acrylic and protected with foil-lined conical shields.

### Training Procedures

Behavioral training was performed as previously described (Lee et al., 2021). Briefly, mice were water-deprived to 85% of baseline following post-operative recovery. Mice were habituated to head fixation inside the custom sound-proof chamber for 1-4 days. Licks were detected with an infrared emitter and sensor positioned at the sucrose/quinine lick spout. Tasks were automated with custom LabView scripts on a laptop connected to a DAQ port (National Instruments). Mice were trained to lick actively for sucrose in response to an LED cue following a non-odorized air puff (1-5 days). They were then trained to withhold licking during the 2 s delay period between air and LED cue (1-5 days). For pre-learning of the go/no-go task, mice learned to lick for sucrose in response to Go Odor-A (isoamyl acetate) and withhold licking to No-Go Odor B (α-pinene) to avoid quinine. Pre-learning typically took 5-10 days.

*Associative learning.* A ∼20-trial session with random trials of only Odor-A and Odor-B was tested first (AB-only session). Immediately afterwards, two novel odors in addition to Odor-A and Odor-B were presented in random order every 18-22 s (AB12 session). One of the novel odors was associated with lick, and the other with no-lick. Mice learned this new association by trial and error. Novel odor pairs were never repeated for an individual mouse. The lick-associated odors (Odor-C, -E, -G, …) were collectively termed as Odor- 1, and the no-lick-associated odors (Odor-D, -F, -H,…) were termed as Odor-2. Odor-A and Odor-B were randomly delivered with 20% emergence rate each, whereas Odor-1 and Odor-2 were delivered with 30% emergence rate each. Licking during the delay period automatically aborted the trial. Odors used in the study are summarized in **Extended Data Table 3**.

### Data Collection

*Spike recording.* Spiked data was acquired as previously described (Lee et al., 2021). Tetrodes were advanced 40-80 µm to record a different group of cells in each session of a same type (either control or inhibition session) at the same recording position, assuring that identical cells were not double sampled. We typically advanced tetrodes after the recording of one session.

*Optogenetic Stimulation.* Laser (473 nm, 20–25 mW) was only held on from the Odor-1 cue onset for 5 s, followed by a linear taper of 10 s on each trial. For control experiments, we used no-laser sessions in the same mice used for stimulation, where disconnected laser patch fiber tips were placed inside the implanted conical shield and laser was applied inside the shield with the same condition as stimulation sessions. Our previous experiments validated that this inter-animal comparison allows us to reduce variability between individuals, and has a similar to better statistical power compared to the usage of non-opsin GFP control groups in different animals (Lee et al., 2021). To control for the unequal number of sessions from individual mice, we averaged behavioral performance across sessions for each mouse and used these mean values (see Data Analysis below).

*L-3,4-dihydroxyphenylalanine (L-DOPA) treatment*. L-DOPA (5 mg/ml in saline, 10 ml per kg body weight) was injected intraperitoneally into APP-KI mice 30 minutes before the behavioral test (L-DOPA group). For mice in the control group, saline (10 ml per kg body weight) was instead injected. Injection was carried out for five consecutive days, and behavioral data during these five days were used for the analysis.

*Photometry recording*. The photometry recording was performed using a custom-made photometry setup as described previously (Lee et al., 2021). In brief, the excitation laser (473 nm, 0.4–0.5 mW) was delivered through a dichroic and a patch cord (Doric Lenses). Activity-dependent fluorescence emitted by GCaMP-expressing axons was spectrally separated from the excitation light using a dichroic, passed through a single-band filter, and focused onto a photodetector connected to a current preamplifier (SR570, Stanford Research Systems). The output voltage signals from the preamplifiers were collected by an AD board (National Instruments).

### Data Analysis

#### Unless indicated otherwise, analyses were performed using MATLAB codes written by the authors

*Behavioral analysis*. Behavioral performance was calculated as previously described (Lee et al., 2021). Briefly, instantaneous behavioral performance for Odors-A/B and Odors-1/2 was calculated within a sliding window of 20 trials. Percent correct trials in T5 was calculated from trials 121-160. To evaluate the behavioral performance for each mouse, these two values were averaged for all sessions for each mouse. “Correct” sessions were considered as those with (percent correct trials in T5) > 80%.

*Spike sorting.* Spike sorting was performed as previously described (Jun et al., 2021; Lee et al., 2021). Putative principal neurons with spike peak-trough width of more than 230 µs and cells with more than 100 spikes in a session were included for further analyses. For Figure S4, all neurons were analyzed. Double-counted cells were removed by comparing clusters before and after tetrode turns.

*Spike response.* Mean firing rates in 50-ms bins were obtained using a Gaussian filter with a sigma of 100 ms and transformed as z-score using the mean firing rate during the baseline pre-odor period (-1 – 0 s from odor onset). Significant response was assessed from total spike numbers during odor (0.5 – 1.5 s after odor onset), delay (2 – 3 s after odor onset), and choice (3 – 4 s after odor onset) periods, compared with those in 1 s pre-odor period (Wilcoxon signed-rank test). We used 0.5 – 1.5 s after odor onset for the odor response period because of the delay from odor delivery to the onset of LEC activity observed in this and previous studies (Igarashi et al., 2014; Lee et al., 2021).

*Principal component analysis (PCA)*. Principal component analysis was performed as previously described (Lee et al., 2021). Principal components (PCs) 1 and 2 were used for the subsequent analyses. Euclidian distances were calculated between each odor trial type. The Euclidian distance during the odor period of 0.5 – 1.5 s after odor onset was assessed as mean distance between odor trial types.

*Shuffle analysis.* Shuffle analysis was used to statistically evaluate the mean distances between odor types (Igarashi et al., 2014; Lee et al., 2021). Shuffle data were obtained by randomly shuffling the assignment of odor type for each trial while keeping the total number of each trial type same as the original real data. With this shuffling procedure, the distinct response to specific odor types observed in the original data disappeared, producing randomly distributed spike responses across each odor trial type. Mean distance during the 1.5 s period of 0.5 – 1.5 s after odor onset was obtained for each shuffled data. This procedure was repeated 1000 times, producing 1000 distances for each odor pair. The upper 95^th^ percentile of 1000 distances from each odor pair was averaged and used as a threshold for the statistical assessment.

*Similarity index.* A mean shuffle distance was obtained by averaging the shuffle distance across T1 – T5. A similarity index (SI) was calculated as the difference between real and mean shuffled distance, normalized by the mean shuffled distance (Lee et al., 2021):

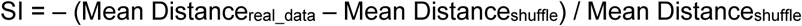

The numerator of SI was negatively flipped so that SI becomes positive if the distance obtained from a given odor pair was smaller than shuffle distance (similar representations of two odors), and negative if the distance was larger than shuffle distance (dissimilar representations of two odors).

*Bootstrapping analysis.* The change of SI during associative learning was compared using the bootstrapping method as previously described (Lee et al., 2021). Briefly, PCA was performed from a resampled neuronal population, and this procedure was repeated 1000 times to make 1000 bootstraps. SI was calculated for each bootstrap, then SIs in T2 – T5 were subtracted by that in T1, to test for a significant distribution above or below zero. Distribution of difference with p<0.05 above or below zero was considered significant. The bootstraps were also compared between WT and APP-KI mice.

*Decoding analysis.* Mean firing rates in 50-ms bins during 1–4 s after odor onset for each odor type (A, B, 1 and 2) were normalized and processed for decoding analysis using binary support vector machine (SVM) learners. Mean firing rates for Odor-A and Odor-B were used to train the SVM. To test performance of prediction, mean firing rates for Odor- 1 and Odor-2 were used. This procedure was repeated 10 times to make 10 bootstraps. For each bootstrap, the decoding performance rate predicting Odor-1 or Odor-2 during the 3-s period from 2-5 s after odor onset was calculated.

### Histology and Reconstruction of Recording Positions

*Electrode positions.* Recording positions were confirmed by passing electrical current through tetrodes in anesthetized mice, followed by fixation in 4% paraformaldehyde and cryosectioning (Jun et al., 2021, Lee et al., 2021). Only data from tetrodes in LEC were collected for analysis. Electrode position in each recording session against the layer 2/3 of the LEC was estimated using following two criteria: (1) extrapolated position from the histological lesioning and tetrode turning history, and (2) numbers of cells obtained in each session. Typically, numbers of cells become maximum when tetrodes hit the LEC layer 2/3. Using this estimated position, we collected neurons ± 200 µm from putative LEC layer 2.

*Immunostaining.* Sections were rinsed three times for 10 min in 1× PBS (pH 7.6) at room temperature, then preincubated for 1 h in 10% normal goat serum in PBST (1× PBS with 0.5% Triton X-100). Between incubation steps, sections were rinsed in PBST. Sections were incubated with antibodies against tyrosine hydroxylase, raised in rabbit (MB152, Millipore, 1:1,000), or amyloid beta, raised in mouse (McSA1, MEDIMABS, 1:500) for 24 h in antibody-blocking buffer at 4 °C. After three 15-min washes in PBST at room temperature, sections were incubated in a goat anti-rabbit antibody conjugated with Alexa Fluor 488nm (ab150077, Abcam, 1:250), or goat anti-mouse antibody conjugated with Alexa Fluor 488nm (A-11001, Thermo Fisher Scientific, 1:250) for 2 h at room temperature. After rinsing in PBS, sections were mounted onto glass slides with 4’, 6’- diamidino-2-phenylindole (DAPI)-containing mounting solution (SouthernBiotech), and a coverslip was applied. Digital photomicrographs were acquired with an Olympus BX53 fluorescence microscope equipped with a digital camera.

*Axon and cell density counting.* For density analysis, images were taken spanning 3.4 –3.8 mm AP from bregma. The LEC is delineated, and the areas of pixels with fluorescence values higher than three standard deviations of background were quantified for all blocks using ImageJ software, and then divided by the whole area of the image for percentage. The density of labelled neurons from the TH+ population in VTA and SNc was evaluated by anti-TH immunostaining. Images were taken from each hemisphere spanning 3.4 –3.6 mm AP from bregma. The number of cells with fluorescence was counted using ImageJ software, and then divided by the whole area of the image for percentage. All analyses were performed blind.

### Statistics and Reproducibility

Data are shown with ± standard error. The animal numbers and sampled neuron numbers (biological replicates) were designed to achieve a power of greater than 0.8. Both sexes of animals, randomized and blinded analyses were used. For statistical testing, data were first tested for normal distribution using the Kolmogorov-Smirnov test (p<0.05 cut-off). All statistical methods used are summarized in Extended Data Table 1 and were two-sided.

## Extended Data Figure Legends

Extended Data Figure 1 | Aβ deposition in APP-KI mice

(a) Coronal section with anti-Aβ immunostaining for 12-month-old WT and 1, 2, 3, 4, 6, 9 and 12-month-old APP-KI mice. D, dorsal, V, ventral, M, medial, L, lateral.

(b) Density of Aβ in the LEC of WT and APP-KI mice across ages. p=1.8e-7, ANOVA; p=2.3e-4 or less, post-hoc Tukey test; n = 4 mice from each age group.

(c) Same as Fig. S1b but for density of Aβ in the VTA (left) and SNc (right) (VTA: p=7.4e-7, ANOVA; p=0.038 or less, post-hoc Tukey test. SNc: p=7.4e-7, ANOVA; p=0.035 or less, post-hoc Tukey test; n = 4 mice from each age group).

Extended Data Figure 2 | Detailed behavior analysis of WT and APP-KI mice

(a) Detailed behavioral performance of young WT and young APP-KI mice in Fig. 1c- d, plotted for each odor trial type. Learning curves during young WT (Left) and young APP-KI (Middle). (Right) Performance of mice in T5 (p=0.021, ANOVA; p=8.1e-6 between Odor-1 in young WT mice vs. Odor-1 in young APP-KI mice, post-hoc Tukey test; n = 11 young WT mice, n= 16 young APP-KI mice). Other p- values not shown in the figure were non-significant.

(b) Same as Extended Data Fig. 2a but for old WT and old APP-KI mice. (p=6.9e-5, ANOVA; p=1.5e-15 between Odor-1 in old WT mice vs. Odor-1 in old APP-KI mice, post-hoc Tukey test; n = 11 old WT mice, n= 14 old APP-KI mice).

(c) Percentage of correct trials in T5 of young WT vs old WT mice (p= 0.031, ANOVA; p= 0.038 between Odor-1/2 in young WT vs. Odor-1/2 in old WT mice, post-hoc Tukey test; n = 11 young WT mice, n = 11 old WT mice).

(d) Percentage of sessions where mice correctly learned new association (p=0.53, binomial test).

(e) Performance of young and old WT mice in T5, plotted for each odor trial type. (p=0.71, ANOVA; n = 11 young WT mice, n= 11 old WT mice).

(f) Days required for obtaining performance over the criteria (80%) during pre-learning sessions in WT and APP-KI mice (p=0.09, ANOVA).

Extended Data Figure 3 | Histological validation of implanted sites

(a) Recording positions in the LEC of WT and APP-KI mice for electrophysiology recording experiments (Fig. 2 and Extended Data Fig. 9, 10). Tetrode tips obtained using electrolytic lesioning were marked with arrowheads. D, dorsal, V, ventral, M, medial, L, lateral.

(b) Optic fiber positions in the LEC of young DAT-Cre and young APP-KI x DAT-Cre mice injected with AAV-flex-GCaMP6m into the VTA and SNc for photometry experiments (Fig. 3). Arrowheads show estimated tips of optic fibers. Four mice received unilateral implantations and eight mice received bilateral implantations.

(c) Optic fiber positions in the LEC of APP-KI x DAT-Cre mice injected with AAV-flex- ChR2-eYFP or AAV-flex-ChR2-mCherry into VTA and SNc for stimulation of LEC-projecting VTA/SNc dopaminergic fibers in LEC (Fig. 4).

Extended Data Figure 4 | Spike properties of LEC_L2/3_ neurons in young WT and young APP-KI mice

(a) Spike properties of LEC_L2/3_ neurons. Top, plot of individual neurons for half amplitude duration as a function of spike width in young WT (left) and young APP-KI (right) mice (n=479 in young WT mice and n=846 in young APP-KI mice). Dashed lines (230 and 390 µs) represent cut offs for classifying three neuron groups: narrow spiking (NS) cells, regular spiking (RS) cells and wide spiking (WS) cells. Middle, distributions of neurons as a function of spike width. Trimodal distribution of spike widths reveals three groups (NS cells, RS cells and WS cells). Bottom, percentage of NS, RS, and WS cells in young WT and young APP-KI mice. (p=1.2e-23, Chi-squared test for comparing distributions; ***p<0.001 for all cell types, Benjamini-Hochberg post-hoc test for comparing each cell type).

(b) Mean spike widths of putative principal cells (NS cells and WS cells) and putative interneurons (NS cells) between young WT and young APP-KI mice (p=3.9e-15 for principal cells, p=0.76 for interneurons; rank sum test).

(c) Mean firing rates of LEC_L2/3_ putative principal cells throughout the associative memory learning session. (c1) Left, mean firing rate (p=3.9e-15, rank sum test). Right, a cumulative distribution plot (p=3.3e-12, Kolmogorov-Smirnov test). (c2)

Left, peak firing rate (p=4.7e-5, rank sum test). Right, a cumulative distribution plot (p=4.4e-4, Kolmogorov-Smirnov test).

(d) Same as Extended Data Fig. 4c, but for LEC_L2/3_ putative interneurons. Mean firing rates, p=0.0044, rank sum test; p=0.031, Kolmogorov-Smirnov test. No significant difference was observed for peak firing rates.

Extended Data Figure 5 | Cue response type of LEC_L2/3_ neurons in young WT and young APP-KI mice during learning

LEC_L2/3_ principal neurons were classified for their cue response type during 0.5-1.5 s after cue onset and shown in percentage among all recorded neurons. Neurons were collected from both correct and error sessions. APP-KI mice had more Odor-A cells and Odor-A/1 cells compared to WT mice through T1-T5. Top, first 10 trials (T1); middle, 10 trials (T3); bottom, last 10 trials (T5) (p=5.5e-4 or less, Chi-squared test for overall distributions; *p<0.05, **p<0.01, Benjamini-Hochberg post-hoc test for each responsive type; n = 423 cells from young WT vs. n= 636 cells from young APP-KI mice).

Extended Data Figure 6 | Selectivity profile of LEC_L2/3_ single neurons in young WT and young APP-KI mice during learning

Example LEC_L2/3_ neurons from young WT(a-d) and young APP-KI mice (e-h). Two neurons were shown each for correct and error sessions. APP-KI mice tended to have more Odor-A cells and Odor-1 cells responding to a single odor cue. *p<0.05, **p<0.01, ***p<0.001 during cue-delay period, Wilcoxon signed-rank test. Black trace, first 10 trials. Red trace, last 10 trials.

Extended Data Figure 7 | Principal component analyses of LEC_L2/3_ neurons of young WT and young APP-KI mice through T1 – T5

(a) PCA trajectories of neural firing of LEC_L2/3_ cell population in young WT mice as in Fig. 2d, 2g, 2h but presented throughout timepoints T1 – T5.

(b) PCA trajectories of neural firing of LEC_L2/3_ cell population in young APP-KI mice as in Fig. 2f, 2g, 2h but presented throughout timepoints T1 – T5.

Extended Data Figure 8 | | Principal component analyses and decoding analyses of LEC_L2/3_ neurons separated for correct and error sessions

(a) PCA trajectories (left), Similarity Index (right top) from spike activity of LEC_L2/3_ population in young WT mice.

(b) Decoding analysis where a support vector machine (SVM) was trained using activities from Odor-A vs Odor-B and tested for performance to discriminate between Odor-1 vs. Odor-2. Bootstrapping method (n=10) was used to obtain results from resampled data. Decoding performance from correct sessions and error sessions were compared using ANOVA (p=6.1e-34, ANOVA; p=2.1e-5 or less between correct session vs. error session in young WT mice at each time point, post-hoc Tukey test). The decoding analysis showed that neural activity in correct sessions discriminated between Odor-1 vs. Odor-2, whereas activity in error session could not discriminate them in WT mice.

(c) Same as Extended Data Fig. 8a, but for spike activity in young APP-KI mice.

(d) The decoding analysis from APP-KI mice showed that neural activity in correct sessions discriminated between Odor-1 vs. Odor-2, while activity in error session could not discriminate them also in APP-KI mice (p=3.7e-21, ANOVA; p=2.8e-4 or less between correct session vs. error session in young APP-KI mice at T2, T3, T4, T5, post-hoc Tukey test).

Extended Data Figure 9 | | Disrupted cue-reward association of LEC_L2/3_ neurons in old APP-KI mice

(a) An example LEC_L2/3_ cell from an old WT mouse showing constant firing for Odor- A and Odor-1. (*p<0.05, **p<0.01 during cue-delay period, Wilcoxon signed-rank test). Black trace, first 10 trials. Red trace, last 10 trials.

(b) An example LEC_L2/3_ cell from an old APP-KI mouse exhibited constant firing only for Odor-1.

(c) (Top) Spike firing rate of n = 671 LEC_L2/3_ cells from old WT mice, shown in z-score during first 10 (T1), middle 10 (T3) and last 10 (T5) trials of the session. (Middle) Time-resolved PCA trajectories of LEC_L2/3_ cell activity for each odor type (▷ cue- onset; ⋆ cue-offset/delay-onset; □ delay-offset). (Bottom) Euclidian distance between odor types.

(d) Same as Extended Data Fig. 9c, but for old APP-KI mice (n = 618 cells). Trajectories for Odor-A and Odor-1 kept far apart through T1 to T5.

(e) Mean Euclidian distance between each odor type during 0.5 – 1.5 s after cue onset in old WT (left) and old APP-KI mice (right). Ninety-fifth percentile distance obtained from shuffled data denotes significant distance (red line).

(f) Similarity index (SI) between Odor-A and -1 in old WT and old APP-KI mice across learning. Positive SI denotes similarity of representations between Odor-A and Odor-1, whereas negative SI denotes dissimilarity.

(g) Similarity Index (SI) of old WT and old APP-KI mice was compared using bootstrapping method. SI was calculated for 1000 bootstraps, then SIs for old WT were subtracted by SIs for old APP-KI. The subtraction confirms higher similarity in the representations between Odor-A and Odor-1 in old WT mice compared to that in old APP-KI mice (*p<0.05, **p<0.01***p<0.001; n=1000 bootstrapping test).

Extended Data Figure 10 | Principal component analyses of LEC_L2/3_ neurons of old WT and old APP-KI mice through T1 – T5

(a) PCA trajectories of neural firing of LEC_L2/3_ cell population in old WT mice as in Extended Data Fig. 9c, 9g, 9i but presented throughout timepoints T1 – T5.

(b) PCA trajectories of neural firing of LEC_L2/3_ cell population in old APP-KI mice as in Extended Data Fig. 9c, 9g, 9i but presented throughout timepoints T1 – T5.

Extended Data Figure 11 | Density of dopaminergic cell bodies in the VTA and SNc did not differ between WT and APP-KI mice

(a) (From left to right) Coronal sections of the VTA and SNc with anti-Tyrosine hydroxylase immunostaining for young WT, young APP-KI, old WT, and old APP-KI mice. D, dorsal, V, ventral, M, medial, L, lateral.

(b) Number of TH positive cells in the VTA of WT and APP-KI mice (p= 0.03, ANOVA). P-values for post-hoc Tukey test are shown in the figure (n = 4 each from WT mice and APP-KI mice). All post-hoc p-values, including those not shown in the figure, were above 0.05.

(c) Same as Fig. S11B but for number of TH positive cells in the SNc. (p= 0.014, ANOVA). All post-hoc Tukey test p-values, including those not shown in the figure, were above 0.05.

Extended Data Figure 12 | LEC dopamine signals in correct and error sessions

(a) Same as Fig. 3g, but for data from young WT mice separated for correct (black) and error (orange) sessions.

(b) Same as Fig. 3g, but for data from young APP-KI mice separated for correct (black) and error (orange) sessions.

(c) Mean GCaMP signals during 1-4 s after cue onset, plotted separately for correct sessions of young WT and APP-KI mice (top), and error sessions of young WT and APP-KI mice (bottom). Odor-1; p= 0.0012, ANOVA; p= 0.0048 in T1, post-hoc Tukey test; n = 10 and n = 8 hemispheres from WT and APP-KI mice, respectively.

(d) Correlation between behavioral performance and photometry signals for Odor-1. Y-axis shows the trial number after which mice started five consecutive hit trials for Odor-1. X-axis denotes the trial number after which five consecutive trials exhibited GCaMP signals > 3 times standard deviation (SD). Note this 3SD criterion removed some sessions with low GCaMP signals. Each dot represents a single session from young APP-KI mice (p=2.5e-5, Pearson correlation; n = 21 sessions form n = 8 young APP-KI mice.

Extended Data Figure 13 | Detailed behavior analysis for optogenetic LEC dopamine stimulation and L-DOPA treatment

(a) Detailed behavioral performance for dopamine fiber optogenetic stimulation in young APP-KI x DAT-Cre mice in Fig. 4b-d, but plotted for each odor trial type. Learning curves during control (left) and stimulation (middle) sessions. (Right) Performance of mice in the last 10 trials (p=2.9e-10, ANOVA; p=0.0034 or less, post-hoc Tukey test; n = 11 mice). Other p-values not shown in the figure were non-significant.

(b) Same as Extended Data Fig. 13a but for L-DOPA treatment in young APP-KI mice. (p=7.1e-10, ANOVA; p=2.7e-7 or less, post-hoc Tukey test; n = 8 mice for saline, n=9 for L-DOPA). Other p-values not shown in the figure were non-significant.

