## Supplemental Figures for "Early disruption of entorhinal dopamine in a knock-in model of Alzheimer’s disease"

### Extended Data Figure 1

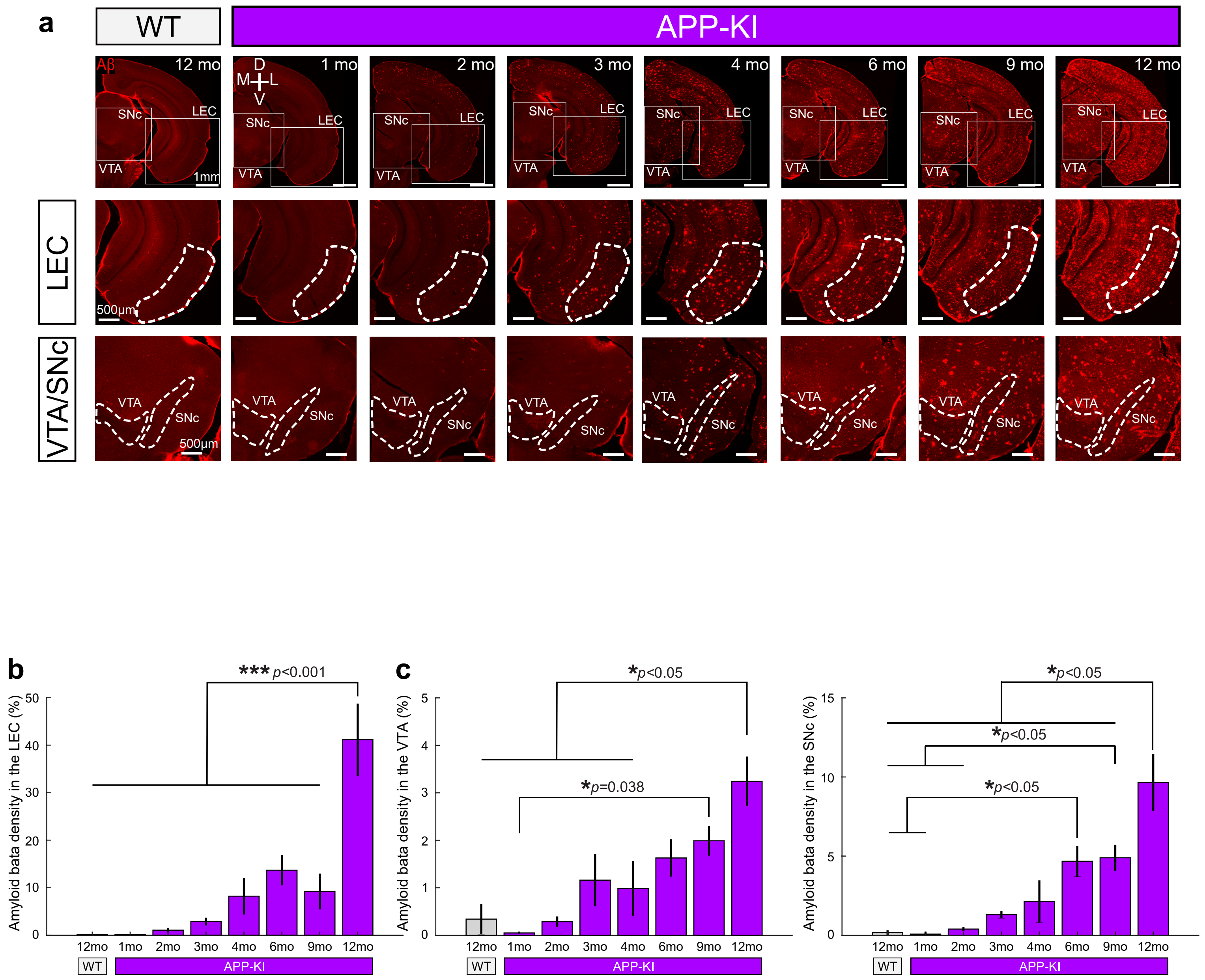

### Extended Data Figure 2

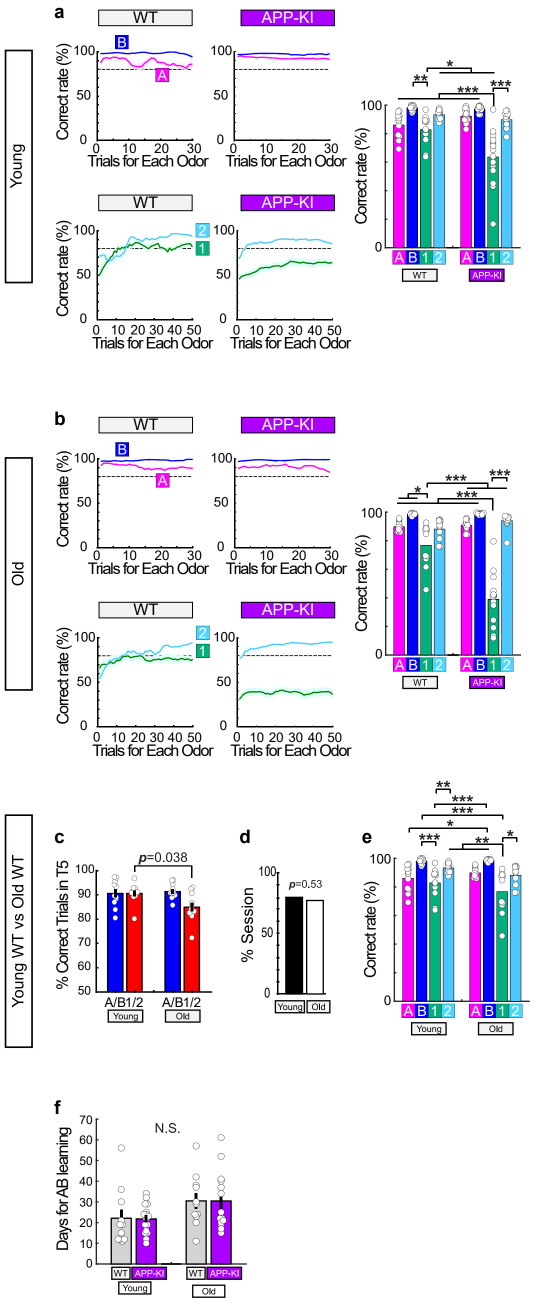

### Extended Data Figure 3

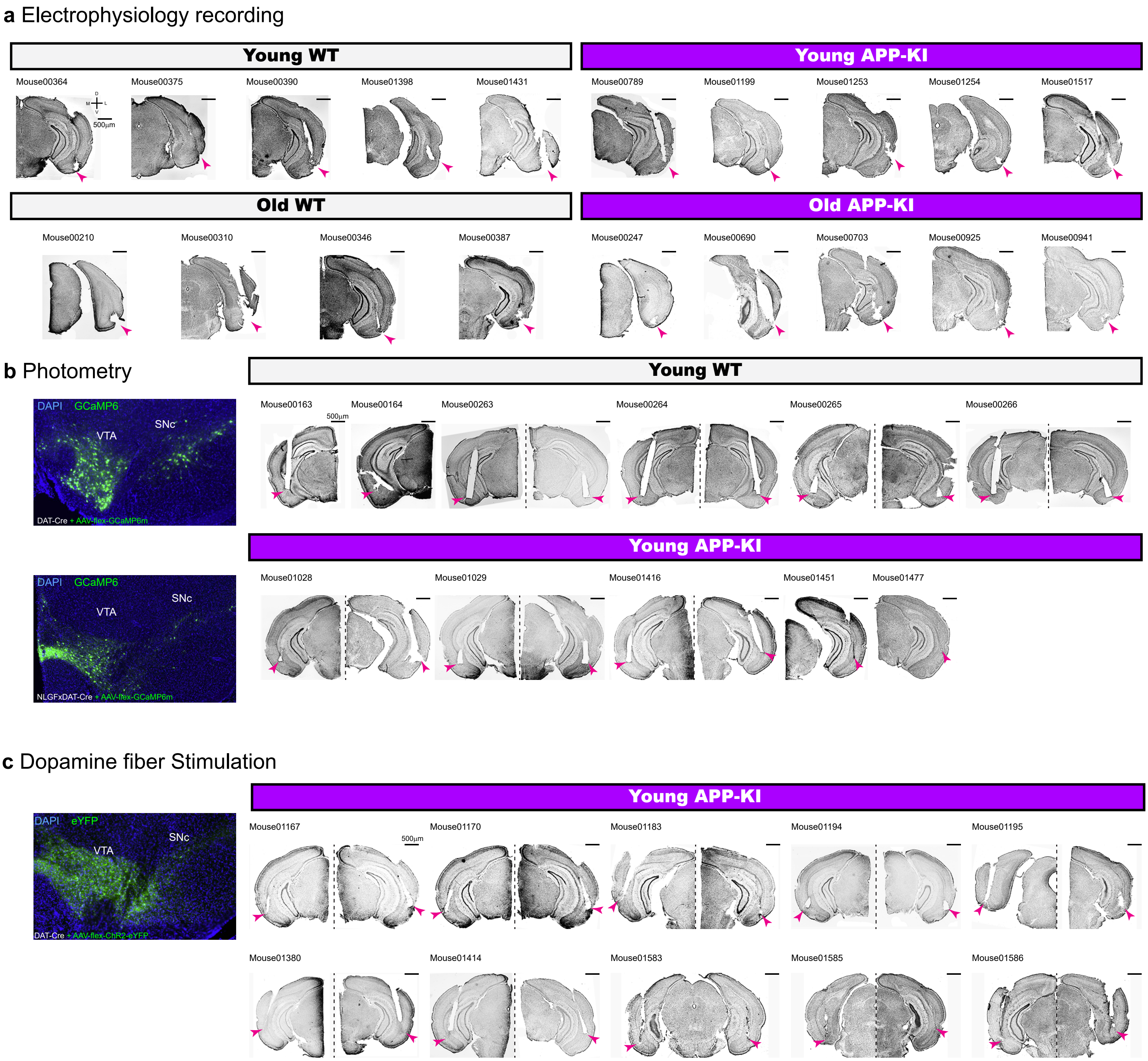

### Extended Data Figure 4

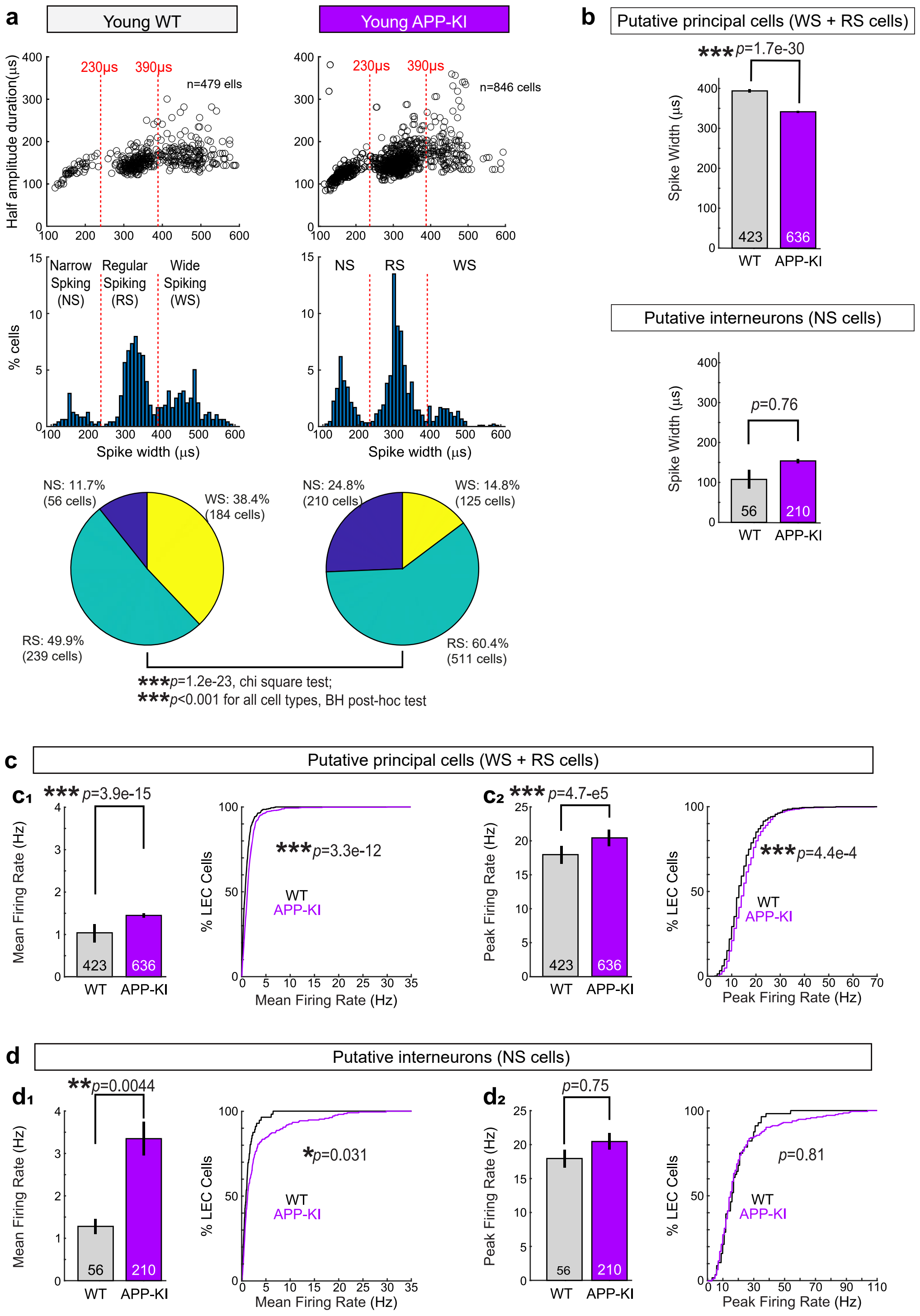

### Extended Data Figure 5

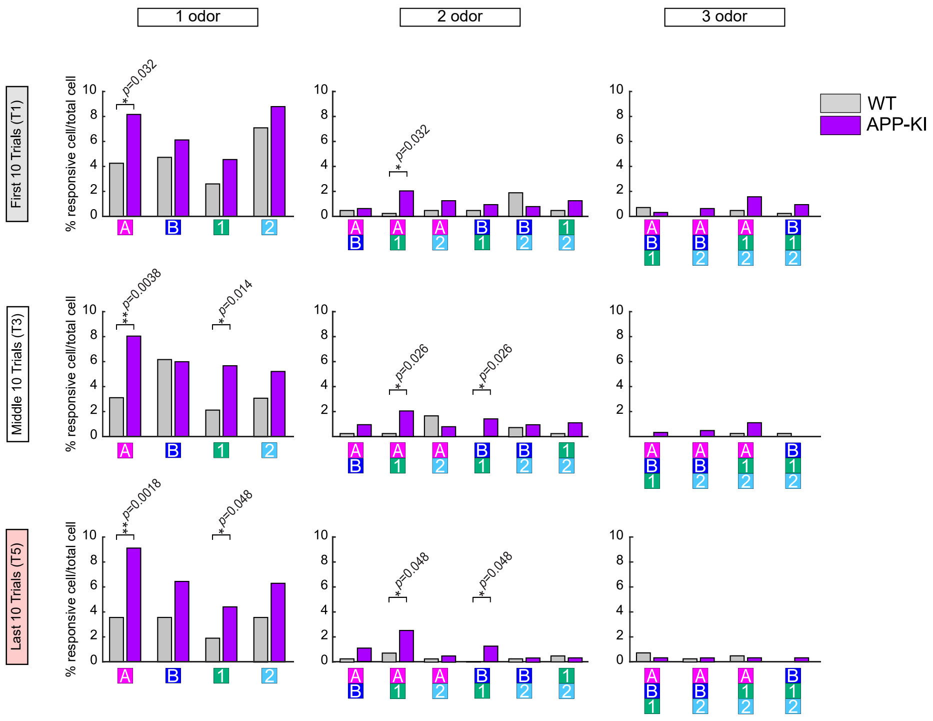

### Extended Data Figure 6

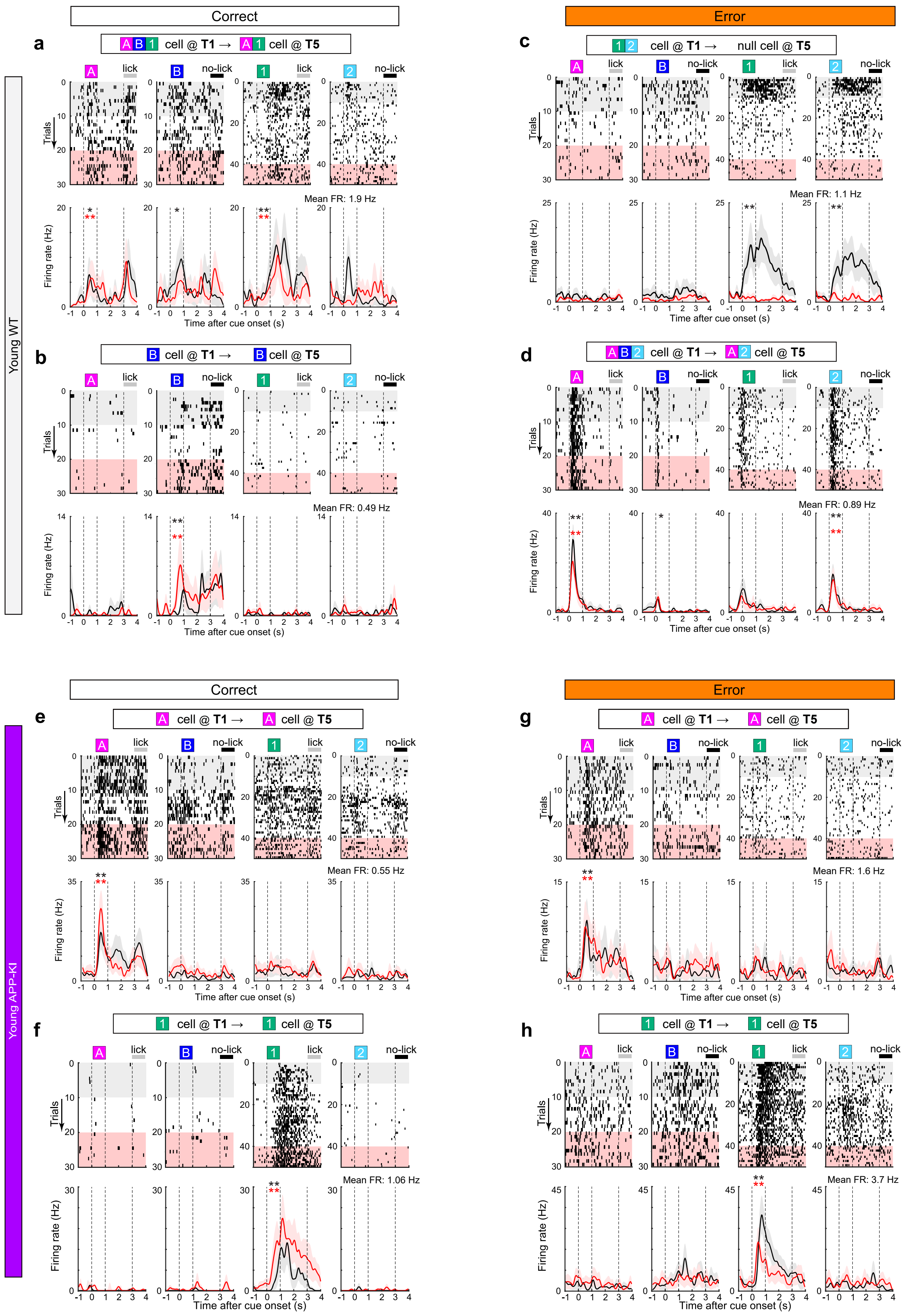

### Extended Data Figure 7

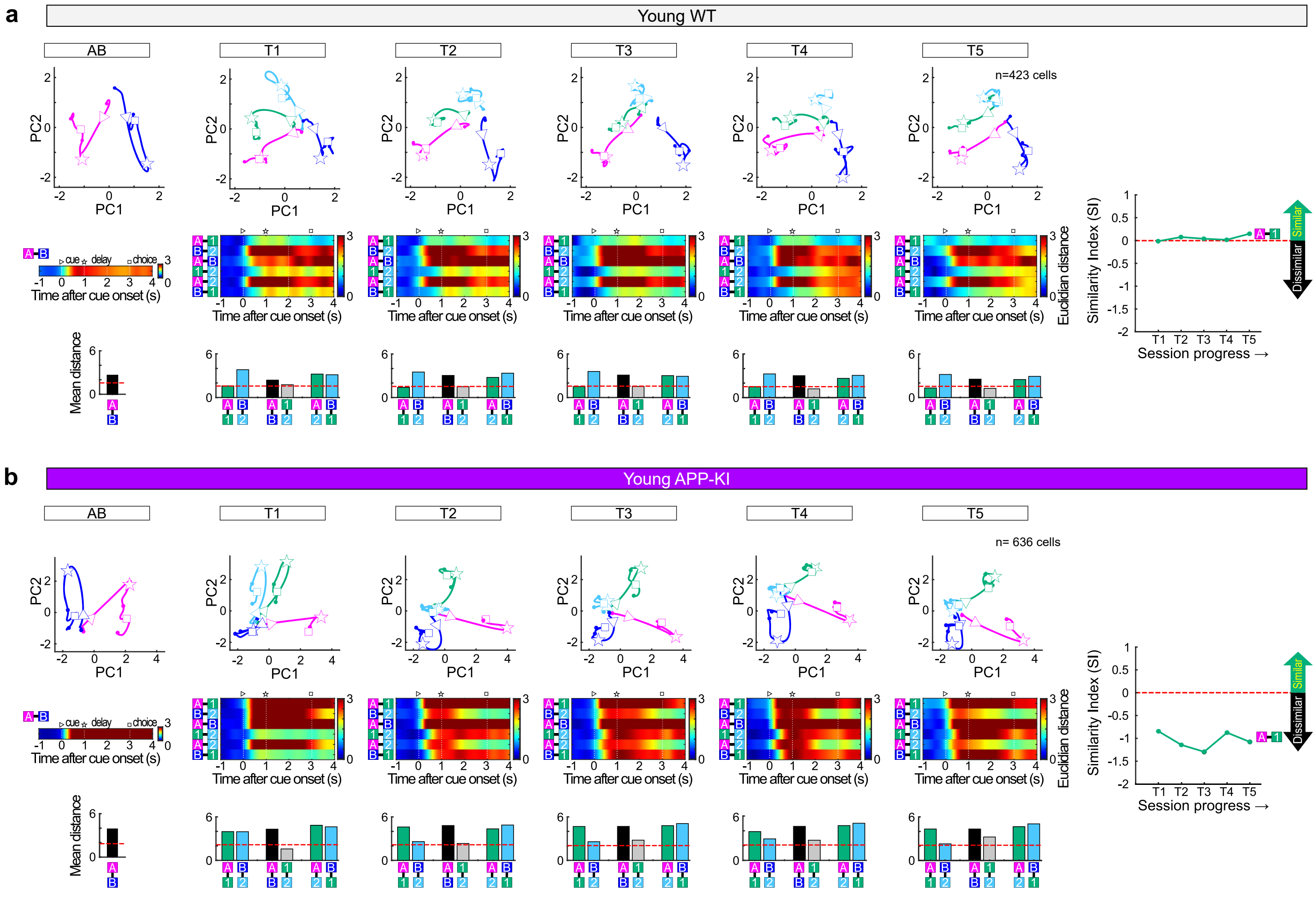

### Extended Data Figure 8

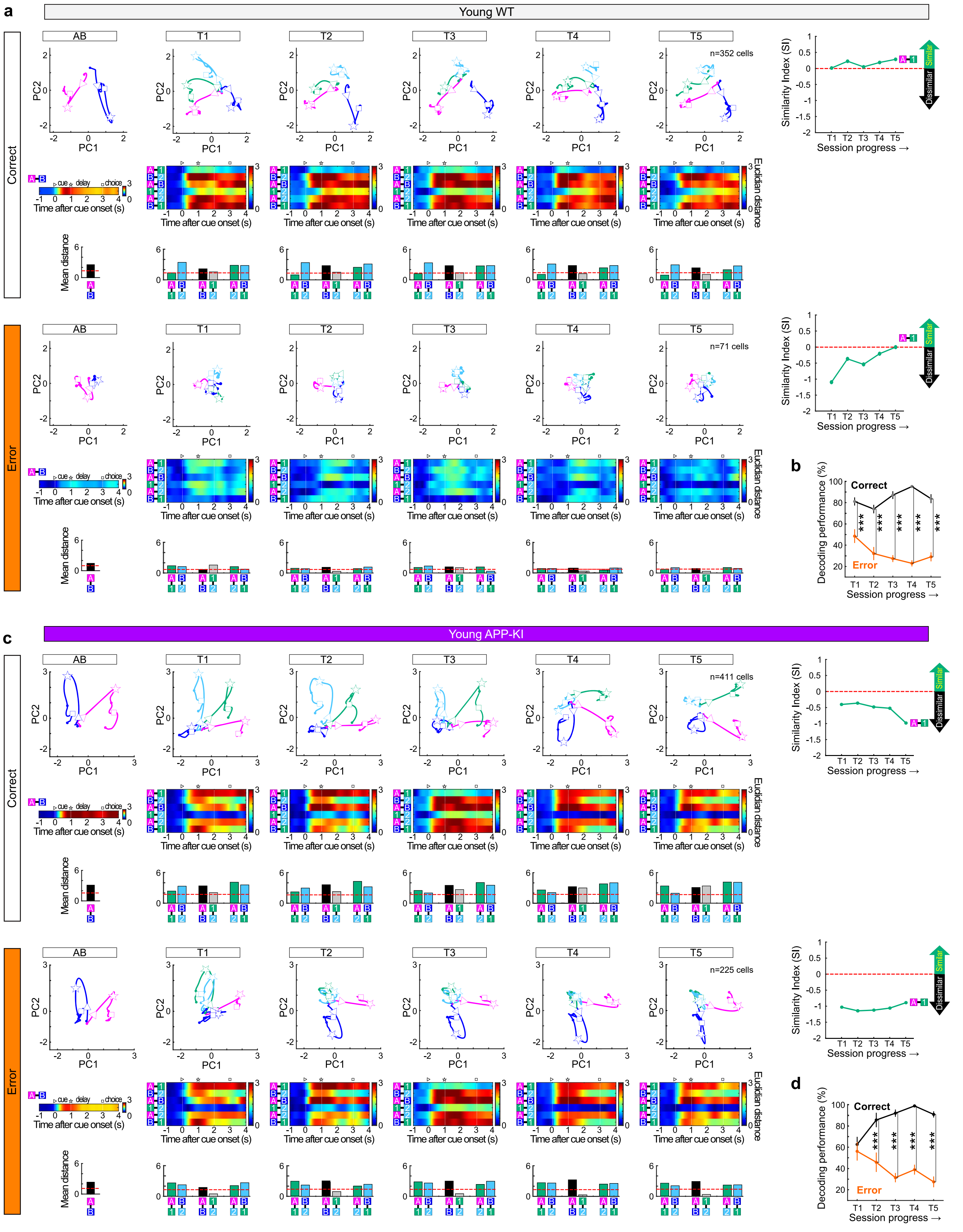

### Extended Data Figure 9

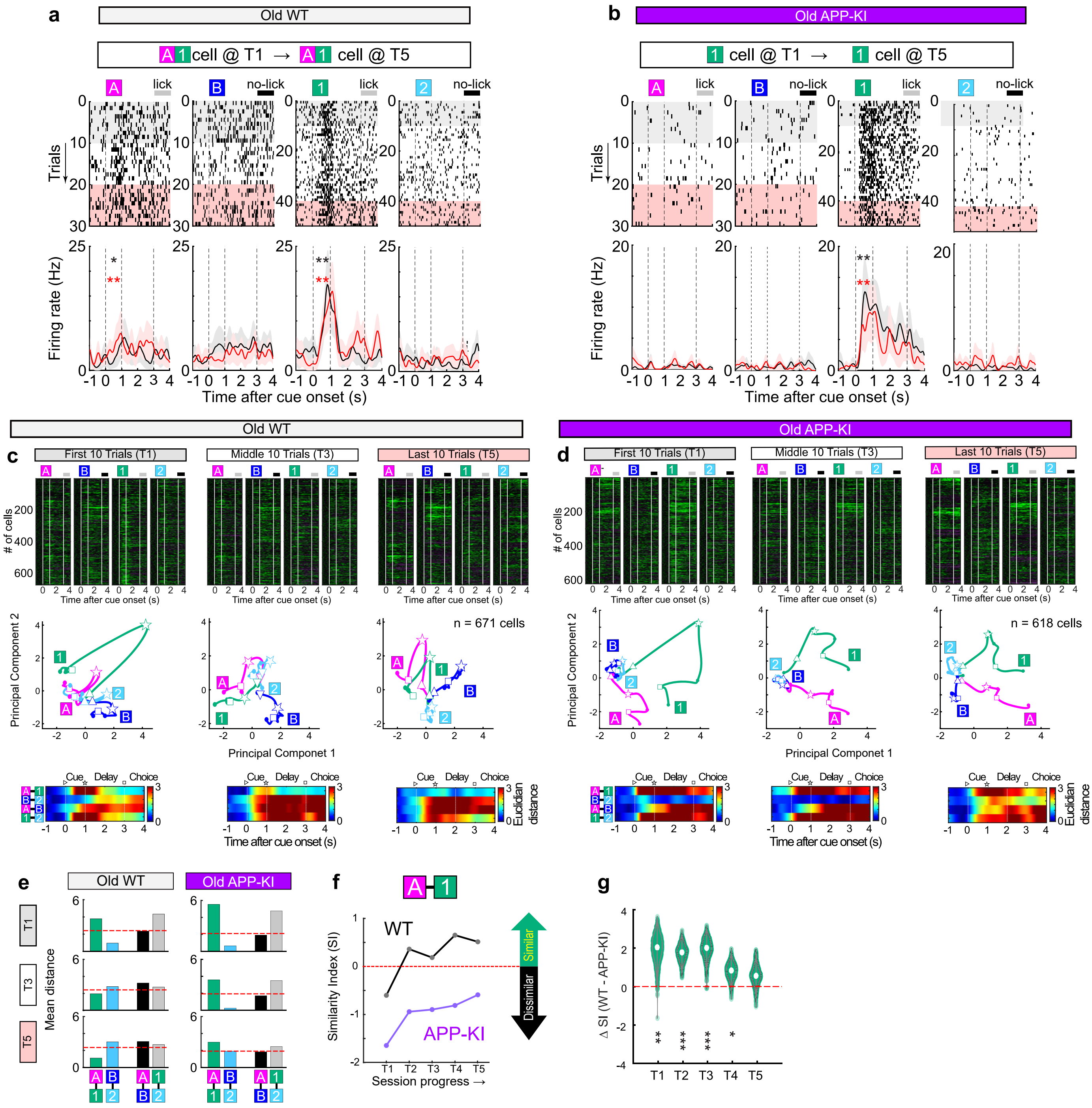

### Extended Data Figure 10

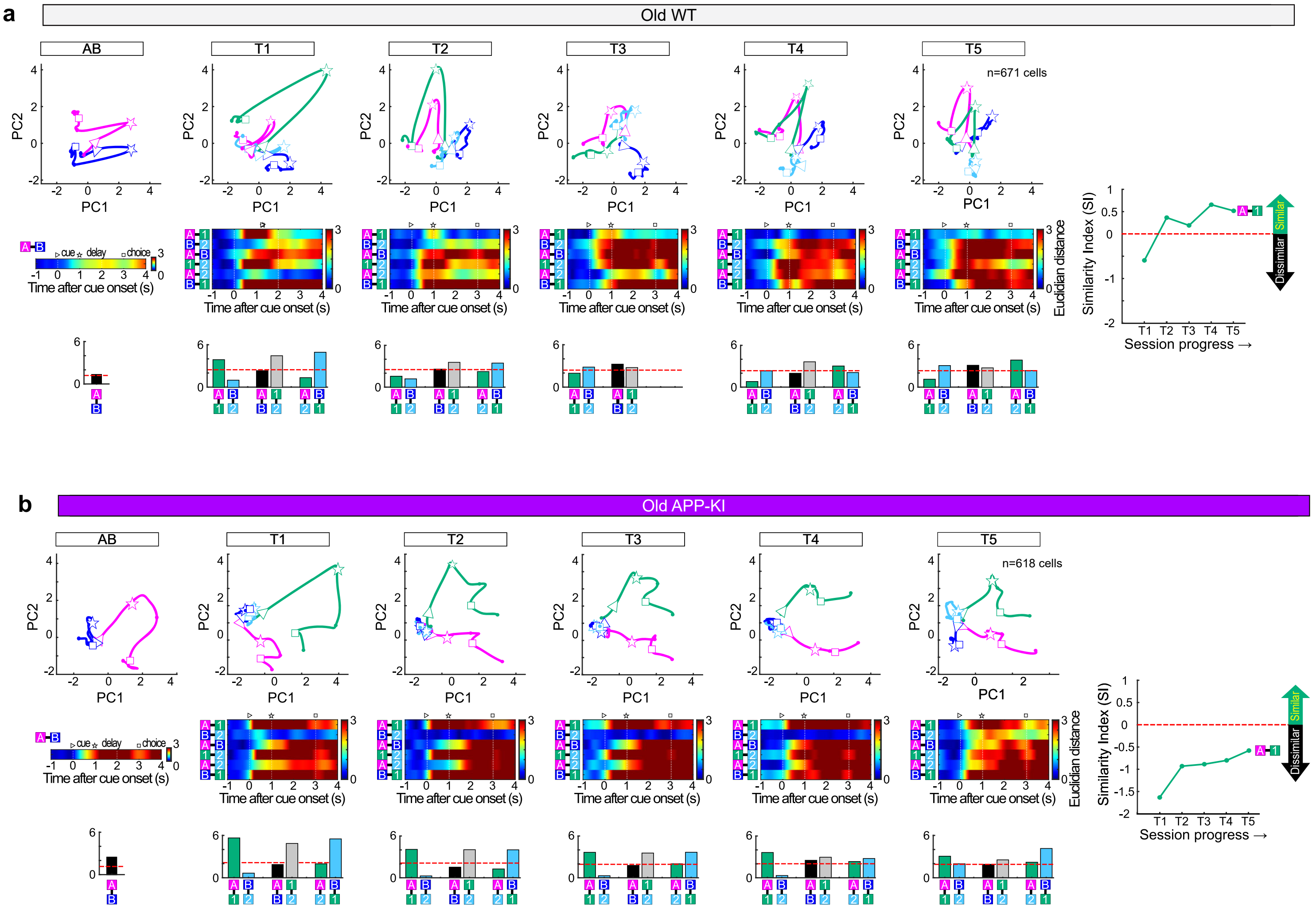

### Extended Data Figure 11

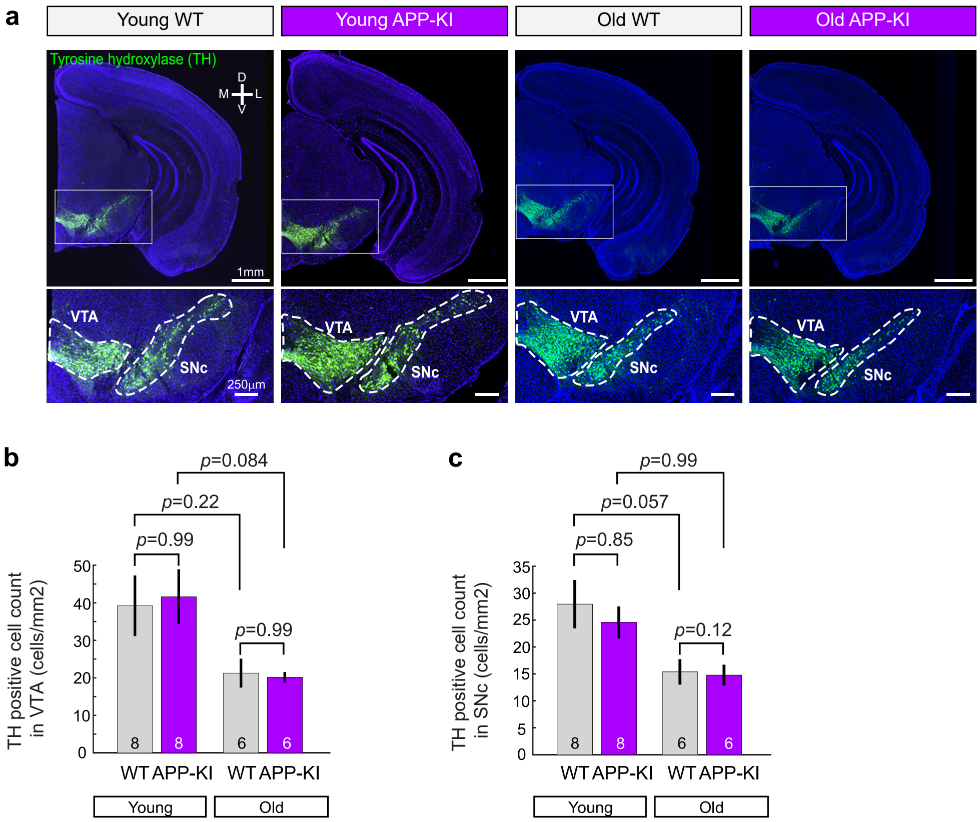

### Extended Data Figure 12

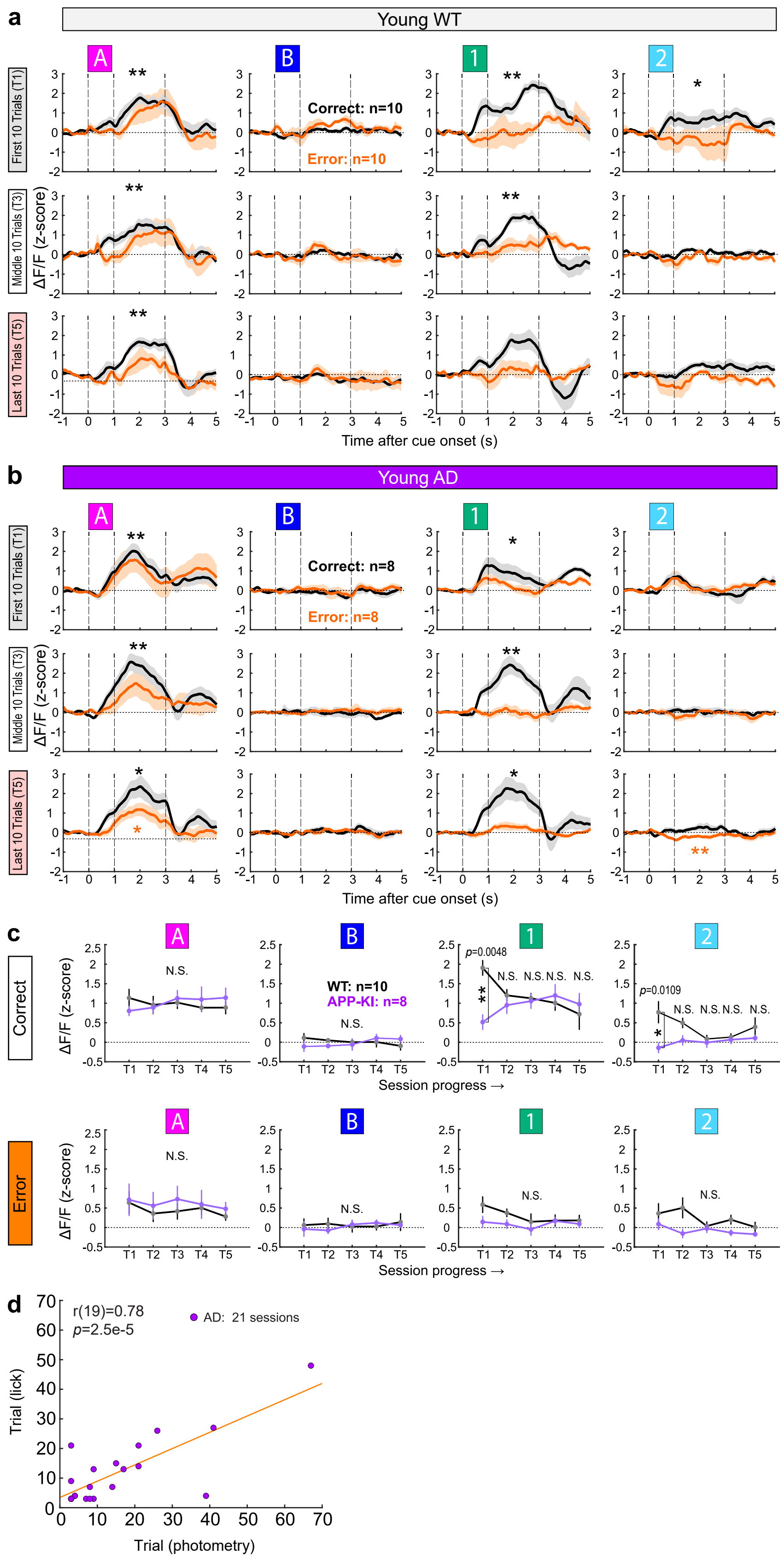

### Extended Data Figure 13

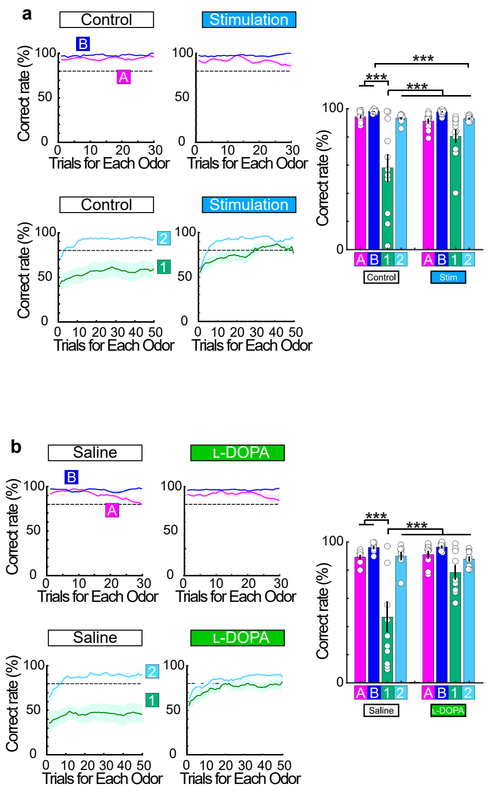
